# Functional Characterization of a Novel Long Non-Coding RNA in *Leishmania braziliensis* Identified Through Computational Screening for Conserved RNA Structures

**DOI:** 10.1101/2025.01.14.632970

**Authors:** Caroline R. Espada, Christian Anthon, Rubens D.M. Magalhães, José Carlos Quilles Junior, Natalia M.M. Teles, Fabiano Pais, Lissur A. Orsine, Letícia de Almeida, Tânia P. A. Defina, Adam Dowle, Jan Gorodkin, Pegine B. Walrad, Angela K. Cruz

## Abstract

*Leishmania* parasites alternate between hosts, facing environmental changes that demand rapid gene expression adaptation. Lacking canonical RNA polymerase II promoters, transcription in these eukaryotes is polycistronic, with gene regulation occurring post-transcriptionally. Although non-coding RNAs (ncRNAs) have been identified in *Leishmania* transcriptomes, their functions remain unclear. Recognizing RNA structure’s importance, we performed a genome-wide alignment of *L. braziliensis* and related species, identifying conserved RNA structures, 38 of which overlap with known ncRNAs. One such ncRNA, *lncRNA45*, was functionally characterized. Using a knockout cell line, we demonstrated that *lncRNA45* is crucial for parasite fitness. Reintroducing the wild-type *lncRNA45* restored fitness, while a version with a single nucleotide substitution in the structured region did not. This mutation also altered RNA-protein interactions. These findings suggest that *lncRNA45*’s regulatory role and protein interactions rely on its secondary structure. This study highlights the significance of structured lncRNAs in *Leishmania* biology and their potential as therapeutic targets. Further research into these ncRNAs could uncover new parasite regulation mechanisms and inspire novel treatment strategies.

## INTRODUCTION

*Leishmania* is a protozoan parasite and the causative agent of a group of diseases collectively known as leishmaniasis (1). Approximately 20 species are pathogenic to humans, causing various clinical syndromes ranging from skin lesions to potentially fatal visceral leishmaniasis (1). *Leishmania (Viannia) braziliensis* is among the most relevant pathogenic species in South America, causing one of the most severe forms of the disease: mucocutaneous leishmaniasis, which can lead to extensive destruction of mucosal tissues, primarily affecting the upper respiratory tract and facial areas (2). There are currently no vaccines for humans, and treatment relies on a limited number of drugs that often have severe side effects and decreasing efficacy due to parasite resistance (2,3).

Unlike other eukaryotes, trypanosomatids such as *Leishmania* possess a unique genetic organization (4,5). These organisms lack canonical RNA polymerase II (RNA Pol II) promoters for individual genes. Instead, Transcription Start Sites (TSS) drive tandem arrays of multiple genes. This results in polycistronic transcription, where functionally unrelated genes within the same polycistronic transcriptional units (PTUs) are transcribed together as a single polycistron (4,6). Each PTU may contain over 200 genes that undergo *trans*-splicing during RNA processing. *Trans*-splicing involves the insertion of a 39-nucleotide capped exon (spliced leader RNA, SL-RNA) at the 5′-end of each mRNA, followed by polyadenylation of the 3′-end of the preceding mRNA (7). Mature mRNAs are then exported to the cytoplasm for translation into proteins. This makes *Leishmania* parasites an exceptional model for post-transcriptional gene expression regulation.

To complete its life cycle, the heteroxenous *Leishmania* must adapt to significant environmental changes (8). During a blood meal of female Phlebotomine sandflies on an infected mammalian host, phagocytes containing intracellular amastigotes (AMA) are ingested. These amastigotes differentiate into procyclic promastigotes (PRO) in the insect gut. After replication, PRO migrate to the anterior gut and further differentiate into intermediate forms, with metacyclic promastigotes (META) being the infective stage. During the next blood meal, META are regurgitated into the dermis of a vertebrate host, where they are phagocytized by macrophages and differentiate back into AMA (8). Therefore, agile regulation of gene expression is essential for an effective transmission and consequently, parasite survival.

Despite the distinctive genetic organization of *Leishmania*, the mechanisms governing gene expression regulation are not fully understood. Post-transcriptional events play a crucial role, including mRNA processing, stability, nuclear-cytoplasmic transport, and translation (6,9,10). RNA-binding proteins (RBPs) are pivotal in recognizing specific sequences or structural motifs in mRNA untranslated regions (UTRs), influencing their stability, localization, or translation (4,5,11).

Non-coding RNAs (ncRNAs) have been shown to act as gene expression regulators in trypanosomatids (12,13). These RNAs are transcribed but not translated into proteins and are arbitrarily categorized as short (<200bp, sncRNAs) or long (>200bp, lncRNAs) ncRNAs (14), a classification that is currently under revision (15). Despite the high number of ncRNAs identified in the transcriptome of trypanosomatids (12,16–18), their biological functions and biogenesis are not fully elucidated. In *L. braziliensis*, 11,372 putative ncRNAs (3,602 of them being differentially expressed across lifecycle stages) have been identified through RNA-seq and comparative transcriptomics suggesting they could be enrolled in gene expression regulation in this parasite (17).

Many ncRNAs, such as tRNAs, rRNAs and miRNAs, are strongly structured and function through structure. LncRNAs appear not too strongly structured but may contain structural regions from which they function (19). In addition, some lncRNAs were shown to function through structural elements rather than conserved nucleotide sequences, highlighting the importance of secondary structures (20).

Based on the importance of secondary structure for ncRNAs’ role in living systems, using a computational approach we searched for conserved secondary structures of ncRNAs in trypanosomatid genomes (21), particularly in *L. braziliensis* MHOM/BR75/M2903, and identified ncRNAs with conserved secondary structures differentially expressed between morphologies, suggesting their biological relevance. Based on differential expression and genomic localization, we selected one lncRNA with a conserved secondary structure to investigate its putative regulatory function. We found that this lncRNA is implicated in parasite growth in culture by interacting with proteins enrolled in gene expression regulation and cell-cell signaling pathways and that modifications in its secondary structure caused by a single base substitution impair its function in *L. braziliensis*. Overall, understanding the role of ncRNAs, particularly their structural dynamics, provides insights into gene regulation mechanisms crucial for the survival and transmission of *Leishmania* parasites.

## MATERIAL AND METHODS

### Quality analysis of Trypanosomatids genome using BUSCO

BUSCO (Benchmarking Universal Single-Copy Orthologs) provides a methodology to quantify the completeness of genomic data sets in terms of the expected gene content based on evolutionary principles (22). BUSCO uses orthologous groups selected from OrthoDB (23), it contains proteins present as single copy orthologs in at least 90% of the species included in a specific phylogenetic group (22,23). Using genome as input sequence, BUSCO software locates candidate regions using local alignment against amino acid BUSCO consensus sequences (24), extracts gene models from these regions based on block profiles (25), and scores candidate genes against the profile Hidden Markov Model (HMM) of corresponding BUSCO genes (22).

BUSCO tool suite (v.4.0.0) metrics was used to assess the quality and completeness of the assemblies of Trypanosomatids genomes. This tool was used in genome mode with lineage specific dataset euglenozoan_odb10 (creation date: 2019-11-21, 31 species, 130 universal single-copy orthologs, https://busco-data.ezlab.org/v4/data/lineages/euglenozoa_odb10.2019-11-21.tar.gz). *Leishmania tarentolae* was used as AUGUSTUS parameter, the only trypanosomatid species with training annotation file available in AUGUSTUS database.

### Genome alignment and search for conserved regions

Local Alignment Search Tool, blastZ-like (Lastz v.1.04.03) was used to align ten trypanosomatids genomes of interest with the *Leishmania braziliensis* MHOM/BR/75/M2903 genome. Lastz was developed specifically to optimize the alignment of large sequences in terms of performance and accuracy (26).

The multiple alignment details from existing text), which covers 99% of the *L. braziliensis* genome is filtered by positive score, 3+ sequences/organisms in the alignment, and at least an alignment size of 20. The after filtering 92% of the *L. braziliensis* genome is covered. The alignment is cut up into windows of size 40-120nt, using step size of 40, and requiring at least 3 sequences in the window. This filters the coverage of the *L. braziliensis* genome by RNAz windows down to 77%. We further require that the windows overlap the lncRNA annotation (details from existing text) and end up with 12,969 windows on which we ran RNAz on both strands.

We obtained a false-discovery-rate for the RNAz prediction by shuffling the alignment windows with SISSIz (27). Each window was shuffled 100 times under conservation of di-nucleotide contents.

### Predictions of mutations that affect the RNA structure

*RNAsnp* tool with parameters Model1 p.value 0.01, Model2 p.value 0.01, winsize 200, winsize extention 200 (28,29) were used to predict point mutations that can affect the stability of RNA secondary structures.

### Cell lines culture

*L. braziliensis* M2903 (MHOM/BR/75/M2903) cell line expressing Cas9 and T7 RNA polymerase from pTB007 plasmid (30) and carrying the reporter gene tdTomato in the SSU locus was used for the experiments (31). Promastigotes were cultured at 25°C in M199 medium (Sigma-Aldrich, St. Louis, MO, USA) supplemented with 2.2 g/L NaHCO_3_, 0.1 mM adenine hemi sulfate, 0.005% haemin, 40 mM 4-(2-hydroxyethyl) piperazine-1-ethanesulfonic acid (HEPES) pH 7.4, 100 µg/mL penicillin/streptomycin (Pen/Strep), 1µM Biopterin and 10% heat-inactivated fetal bovine serum (FBS). The selection drugs were added at 32 µg/ml hygromycin B (Gibco™), 50 µg/ml Nourseothricin (Sigma-Aldrich^®^), and 20 µg/ml puromycin dihydrochloride (Gibco™).

Axenic amastigotes of *L. braziliensis* M2903 were obtained by cultivating metacyclic promastigotes in 100% FBS incubated at 33°C in 5% CO_2_.

THP-1 cells used in infectivity assays were cultured in RPMI media supplemented with 10% FBS and 100 μg/mL Pen/Strep. Cells were kept at 37°C and a 5% CO_2_ atmosphere and passaged after every 3 days or when cell density was close to 10^6^ cell/mL.

### DNA extraction

*L. braziliensis* total DNA extraction for cloning and sequencing purposes was done using DNeasy Blood & Tissue Kits (Qiagen) according to the manufacturer’s specifications. For colony screening the protocol described by (32) was employed.

### RNA extraction

Total RNA was extracted from *L. braziliensis* cultures using the Directzol kit Quick-RNA Miniprep (Zymo Research) according to the manufacturer’s instruction. After extraction, RNA samples were treated with TURBO™ DNAse to remove any DNA remains.

### Real-time quantitative PCR (RT-qPCR)

Differential expression was evaluated by RT-qPCR using primers specific for *lncRNA45* (*lncRNA45*_RT-F and *lncRNA45*_RT-R). The gene coding for 7SL RNA was used as normalizer and amplified using the primers 7SL_RT-F and 7SL_RT-R. For the RT-qPCR reaction, 100 ng of cDNA were incubated with SYBR Green/ROX qPCR MasterMix and 0.6 µM of each primer. The reaction was done using StepOnePlus™ Real-Time PCR System using the program: 95°C for 10 minutes followed by 40 cycles of 95°C for 15 second, 60°C for 60 seconds and 72°C for 20 seconds. The Ct values obtained for each sample were normalized by the 7SL values and differential expression was evaluated relative to the expression in procyclic promastigote stage using the 2ΔΔCt method (33). The list of primers used in this study is presented in Supplementary File 2.

### RNA circularization

The RNA circularization assay was performed as previously described by (34) with some modifications. Initially, 5µg of *L. braziliensis* M2903 total RNA (extracted as described previously) were treated (TAP+) or not (TAP-) with 2µL of Tobacco Acid Pyrophosphatase (FirstChoice® RLM-RACE - Invitrogen™) for 5 minutes at 70°C to promote the decapping of RNAs. The reactions were incubated for 2 minutes on ice and after that 30 unities (U) of T4 RNA ligase (NEB) enzyme were added to the system together with 1x T4 Buffer, 1mM ATP and 40U of RNAse inhibitor RNasin® (Promega) in a final volume of 50µL. The reactions were incubated at 37°C for 2 hours followed by an incubation at 65°C for 15 minutes. Circularizes RNA (cRNA) was then precipitated with 0.1 volumes of sodium acetate (pH 5,2), 2.5 volumes of ice cold 100% EtOH and 2µL de glycogen (20 mg/mL) (Thermo Scientific™) for at least 16 hours. RNA was centrifuged at 16.000 x g for 20 minutes at 4°C, washed with 70% ice cold EtOH and ressuspended in 70uL of nuclease free water. Reverse transcription of cRNA was done using TransScript® II First-Strand cDNA Synthesis SuperMix (TransGen Biotech) according to manufacturer’s instructions with a reverse primer specific for *lncRNA45* positioned 200bp from 5’ end (*lncRNA45*_cRT) and 7uL of cRNA (1ug) template. The cDNA obtained was used as template for PCR reactions using primers positioned 100nt from 5’ (*lncRNA45*_cF1) and from 3’ (*lncRNA45*_cF2) ends of *lncRNA45* and directed to the extremities (Figure 5A). Platinum™ *Taq* DNA Polymerase High Fidelity (Invitrogen™) was used for PCR amplifications in the presence of 5uL of cDNA and of lncRNA_cF1 and lncRNA_cF2 primers at an annealing temperature of 58°C and extension of 3 minutes. PCR product was resolved by electrophoresis and each band were individually purified using the kit NucleoSpin Gel and PCR Clean-up (Macherey-Nagel) according to the manufacturer’s instructions. The obtained DNA was cloned in pGEM®-T Easy Vector (Promega) in the proportion 3:1 (insert: vector) and transformed in *E. coli* DH5-α Cloning was confirmed by PCR using M13 *forward* e M13 *reverse* primers and colonies presenting different fragment sizes had the plasmid DNA analyzed by Sanger sequencing using the same primers. The obtained sequences were trimmed to remove vector sequences and mapped to *L. braziliensis* M2903 genome using Geneious Software. The list of primers used in this study is presented in Supplementary File 2.

### Northern Blotting

Total RNA was extracted from parasites in log phase according to the manufacturer’s instructions using the mirVana™ Total RNA Isolation Kit (ThermoFisher Scientific) and treated with Turbo DNase (ThermoFisher Scientific). All solutions used for RNA samples were prepared in RNA buffer (0.5 M Na₂HPO₄, 1 M H₂NaPO₄, pH 6.9 in RNA-se free water).

To confirm the ncRNA size, 4 µg of total RNA was added to 7.5 µL of glyoxal mix (105 µL of DMSO, 30 µL of deionized glyoxal, and 4.2 µL of RNA buffer) and denatured at 50°C for 40 minutes. Then, 1 µL of loading dye (2% bromophenol blue in RNase-free water) was added per sample and applied to a 2% agarose gel stained with SYBR Green II. Samples were electrophoresed at 150V for approximately 90 minutes, followed by imaging on an iBright imager (Invitrogen^™^). RNA was transferred from the agarose gel to a Hybond®-N+ hybridization membrane (Amersham-Cytiva) overnight by capillarity in SSC buffer (3 M NaCl, 0.3 M tri-sodium citrate, pH 7.0). The membrane was then UV cross-linked at 20,000 µJ/cm² for 1 minute and dried at 80°C for 1 hour.

For radioactive probing, an amplicon generated by PCR from the ncRNA target sequence cloned into a pJET II plasmid was used as a complementary sequence. Amplicons were purified from the PCR reaction and radioactively probed according to the manufacturer’s instructions (DIG Northern Starter Kit - Sigma). The membrane was incubated with the probe in a hybridization buffer (1 g/L tetrasodium pyrophosphate, 5x SSC, 5x Denhardt’s solution (1% Bovine Serum Albumin (BSA), 1% Ficoll, and 1% Polyvinylpyrrolidone (PVP) in water), 0.1% SDS, 100 mg/L heparin) at 42°C for 18 hours, followed by washing three times with 0.1% SDS in 2x SSC. The membrane was then incubated with photographic film at −80°C for seven days. The film was developed and imaged using a conventional camera.

### RNA FISH

Approximately 1×10^7^ cells were harvested by centrifugation at 2000 x g for 10 minutes, washed once with PBS and resuspended in a solution of 4% formaldehyde in PBS. Cells were spread in a glass slide previously treated with 0.01% poly-lysine solution (Sigma-Aldrich^®^) for 30 minutes and washed once with PBS. Type I probes, labeled with Alexa Fluor 546 fluorochrome were acquired from (Thermo Fisher^™^) and hybridization was conducted using ViewRNA™ ISH Cell Assay Kit (Invitrogen™) according to the manufacturer instructions and the protocol described by (35). Cells were imaged in a LSM 780/LSM 7MP Carl Zeiss multiphoton microscope with 0.75 magnification.

### Generation of transfectant lines

#### Transfections

Transfections were performed in 1×10^7^ log-phase promastigotes of *L. braziliensis* M2903 Cas9/T7 using Amaxa^TM^ Nucleofactor^TM^ (Lonza) X-001 program, in a final volume of 160 µL of Tb-BSF 1x (36). Transfected cells were transferred to M199 media for 16h and then cloned in M199-agar plates containing 1µM biopterin and the selection drug. Individual clones were transferred to 24-well plates in M199 media containing the selection drug and screened for the presence of the genomic modification by PCR.

#### Knockout

The cell line lacking *lncRNA45* (Δ*lncRNA45*) was obtained using CRISPR/Cas9 according to the method described by (30) with some modifications. The EuPaGDT (37) repository was used to find PAM (NGG) motifs in the *lncRNA45* region. The 20nt sgRNAs targeting sequence were then chosen based on specificity and to be closest as possible from the 5’ and 3’ ends of *lncRNA45*. The 30nt-long homology arms were chosen to be right upstream (5’) and downstream (3’) Cas9 double strand break point. The PCRs for sgRNA template (lncRNA45_5’sgRNA and lncRNA45_3’sgRNA) and donor DNA generation were done as described previously (lncRNA45_UFP and lncRNA45_DRP) (31,38). After transfection the obtained clones were screened for homozygous replacement of *lncRNA45* by the cassette containing the puromycin-resistance gene using the primers Conf-F and Conf-R which anneals upstream and downstream the region of donor DNA recombination site. The clones in which only the band matching donor DNA size was amplified was considered homozygous and the clones having two bands, one matching *lncRNA45* size and the other matching donor DNA size were considered heterozygous. The list of primers used in this study is presented in Supplementary File 2.

#### Overexpressor

To generate overexpressing cell lines, primers OE_*lncRNA45*_NotI-F annealing 55 bases upstream and OE_*lncRNA45*_BamHI-R annealing 115 bases downstream the predicted sequence of *lncRNA45* were used to amplify the *lncRNA45* region from *L. braziliensis* M2903 genomic DNA extracted from log-phase promastigotes using DNeasy Blood & Tissue Kits (Qiagen). The amplified fragment was purified, digested with *Bam*HI (New England Biolabs) and *Not*I (New England Biolabs) and cloned in pSSU-Sat to generate the plasmid pSSU-OE*lncRNA45*-Sat. The generated constructs (pSSU-OE*lncRNA45*-Sat) was transfected into *L. braziliensis* M2903 log-phase promastigotes and clones were isolated as described before. The presence of the pSSU-OE-*lncRNA45*-Sat in the transfectants was verified by PCR using the primers *lncRNA45*-F and Sat-R. A pSSU-Sat plasmid containing a GFP sequence (pSSU-GFP-Sat) instead of *lncRNA45* sequence was used as control. The presence of pSSU-GFP-Sat in the clones obtained after transfection was verified by PCR using the primers GFP-F and Sat-R. The list of primers used in this study is presented in Supplementary File 2.

#### Add-back

The native sequence of *lncRNA45* (*lncRNA45*WT) and the sequence carrying the C50G mutation (*lncRNA45*MUT) fragments were synthesized by Synbio Technologies. To generate the add-back cell lines *lncRNA45*_NotI-F and *lncRNA45*_BamHI-R were used to amplify the *lncRNA45* WT and MUT versions from the synthetic DNA. The amplified fragment was purified, digested with *Bam*HI (New England Biolabs) and *Not*I (New England Biolabs) and cloned in pSSU-Sat to generate the plasmid pSSU-*lncRNA45*WT-Sat and pSSU-*lncRNA45*MUT-Sat. The pSSU-GFP-Sat plasmid was used as mock control. The generated plasmids were transfected into *L. braziliensis* M2903 Δ*lncRNA45* log-phase promastigotes and clones were isolated as described before. The presence of the pSSU-*lncRNA45*WT-Sat and pSSU-*lncRNA45*MUT-Sat in the obtained clones was verified by PCR using the primers *lncRNA45*-F and Sat-R. The presence of pSSU-GFP-Sat in the clones obtained after transfection was verified by PCR using the primers GFP-F and Sat-R. The list of primers used in this study is presented in Supplementary File 2.

#### HA-tagged cell lines

Target proteins were tagged with an anti-HA epitope using CRISPR/Cas9 as previously described by (30,39). Primers for sgRNA template and donor DNA amplification were obtained by inputting *L. braziliensis* M2903 gene ID in LeishGEdit platform. Primers for confirmation were also retrieved by LeishGEdit. The list of primers used in this study is presented in Supplementary File 2.

### Phenotypic assays

#### Growth curve

Growth rates of the parental and transfectant cell lines were compared by area under the curve (AUC) analysis of a continuous 7-day growth curve. For that, 2 x 10^5^ promastigotes/mL seeded in culture flasks containing M199 medium and incubated at 25°C. The parasite density was estimated by counting every 24h during at least 7 days. The experiment was carried out in biological triplicate and the area under the curve of each strain was determined using the GraphPad Prism 8.0.2 software. The significance was determined by non-parametric t-test when two cell lines were compared or by One-Way ANOVA test followed by Tukey’s multiple comparison test when more than two cell lines were compared.

#### Doubling time

Doubling time was measured as described in (38). Briefly, culture density was adjusted to 1×10^6^ promastigotes/mL in M199. After incubation for 24h at 25°C parasite number was determined by counting. Cell density was adjusted again to 1×10^6^ promastigotes/mL in a new flask. This procedure was repeated for four days and at the end of the doubling time (DT) was calculated.

#### Nutritional stress

Capacity to recover from nutritional stress was compared between the parental and the transfectant cell lines. For this, 1×10^6^ promastigotes were incubated for 4h in the presence of M199 (unstressed) or PBS (stressed). After this period, the plate was centrifuged to remove the supernatant and fresh M199 containing 1mg/mL of MTT [3-(4,5-Dimethyl-2-thiazolyl)-2,5-diphenyl-2H-tetrazolium bromide, Methylthiazolyldiphenyl-tetrazolium bromide] (Sigma-Aldrich) was added to the wells. After 24 hours of incubation, the plate was centrifuged to remove the supernatant and DMSO was added to dissolve the formazan crystals. The absorbance was measured in the spectrophotometer at a wavelength of 570 nm and the percentage of cell viability was calculated in the stressed parasites in relation to the non-stressed parasites for cell line. At least three independent experiments were performed with five technical replicates each. The percentage recovery observed for each transfectant strain was compared to that of WT using the One-Way ANOVA test followed by Tukey’s multiple comparison test using the GraphPad Prism 8.0.2 program.

#### Metacyclogenesis

The promastigotes culture was adjusted to 2.10^5^ promastigotes/ml of M199 and grown for 5 days at 25°C. After this period, the culture was centrifuged at 2000 x g for 10 minutes, the pellet was resuspended in fresh M199 and the initial number of parasites was determined. A gradient of Ficoll 20%, 10% (50% Ficoll 20% + 50% M199) was set up and the parasite suspension was added on top of the gradient (40). The gradient was centrifuged at 1200 x g for 10 minutes with acceleration of 1 and deceleration of 0. The first layer of the gradient containing the metacyclic promastigotes was recovered, washed with PBS and counted. The percentage of metacyclogenesis was determined by the ratio between the total number of promastigotes (initial) and the number of metacyclic promastigotes. The percentage of metacyclics observed for each transfectant was compared to that of the parental cell line by non-parametric Student’s t-test using the GraphPad Prism 8.0.2 program.

#### THP-1 Infection

THP-1 monocytes on the third day of culture (density below 1,106 cells/mL) were plated at a density of 3.10^5^ cells per well in RPMI medium, in the presence of 30 nM of PMA (phorbol-12-myristate-13-acetate) to induce differentiation into macrophages. The cells were incubated for 16h at 37°C in a 5% CO_2_ atmosphere. After this period, the medium containing PMA was removed and replaced with fresh RPMI medium. Cells were incubated for 3 days at 37°C in an atmosphere of 5% CO_2_ for maturation. After this period, metacyclic promastigotes were purified from culture in the stationary phase (5th day) as described herein and used to infect THP-1 macrophages in a 10:1 ratio (promastigotes:macrophage). Since all strains generated and the parental cell line constitutively express the tdTomato reporter protein, the percentage of infected cells and the number of amastigotes per macrophage was determined at 24h and 48h using the ImageXpress Micro XLS System equipment (Molecular Devices, LLC, USA) with the same specifications described by (31). The percentage of infection and the number of amastigotes per macrophage observed for each transfectant strain were compared to the parental line using the One-Way ANOVA test followed by Tukey’s multiple comparison test using the GraphPad Prism 8.0.2 program.

#### Oxidative stress

Capacity to recover from oxidative stress was compared between the parental cell line and the transfectants. Axenic amastigotes were obtained as described in cell culture session. Seventy-two hours before the experiment, axenic amastigotes were transferred from 100% FBS to M199 pH 5.4 medium and cultivated at 33°C in an atmosphere of 5% CO_2_. After this period, 5.10^6^ axenic amastigotes per well were incubated for 24h in the presence (stressed) or absence (non-stressed) of 300mM H_2_O_2_. After this period, the plate was centrifuged to remove the supernatant and fresh M199 pH 5.4, containing 1mg/mL of MTT, was added. After 24 hours of incubation, the plate was centrifuged to remove the supernatant and DMSO was added to dissolve the formazan crystals. The absorbance was measured in the spectrophotometer at a wavelength of 570 nm and the percentage of cell viability was calculated in the stressed parasites in relation to the non-stressed parasites for each strain. At least three independent experiments were performed with five technical replicates each. The percentage recovery observed for each transfectant strain was compared to that of WT using the One-Way ANOVA test followed by Tukey’s multiple comparison test using the GraphPad Prism 8.0.2 program.

### S1m-mediated RNA pull-down

#### In vitro pull-down

In vitro pull-down of *lncRNA45* was done using the protocol described by (41) with some modifications. *LncRNA45* native (*lncRNA45*WT) and mutated (*lncRNA45*MUT) sequences were amplified from synthetic native and mutated *lncRNA45* DNA using the primers *lncRNA45*_PacI-F and *lncRNA45*_NcoI-R. The list of primers used in this study is presented in Supplementary File 2. The PCR product was digested and with *Pac*I and *Nco*I and cloned in the plasmid pUC57-T7-4xS1m. The *lncRNA45* fused to the 4xS1m was in vitro transcribed using the MEGAscript^TM^ T7 transcription kit (Invitrogen) according to the manufacturer’s recommendation. A plasmid containing only the 4xS1m sequence was used for in vitro transcription of the empty 4xS1m control. In vitro transcribed RNA was immobilized in streptavidin-coated magnetic beads (SA-beads) (New England Biolabs). Protein lysate of 10^8^ log-phase *L. braziliensis* M2903 promastigotes was extracted by mechanical lysis using a 29G needle in the presence of SA-RNP-Lysis buffer [20mM Tris-HCl pH 7.5, 150mM NaCl, 1.5mM MgCl2, 2mM DTT, 2mM RNase Inhibitor, 1% Triton X-100 and 1 tablet of cOmplete^TM^ protease inhibitor cocktail (Roche) for each 10mL of buffer] and centrifuged at top speed for 10 minutes at 4°C. After that the protein extract was incubated with 50µL of SA-beads for 8h under agitation at 4°C to remove biotinylated proteins, that would nonspecifically bind to the beads. The supernatant was recovered and incubated with the SA-beads with immobilized RNAs and kept under agitation for overnight at 4°C. After washing four times for five minutes with 700µL of SA-RNP wash buffer [20mM Tris-HCl pH 7.5, 300mM NaCl, 5mM MgCl2, 2mM DTT, 2mM RNase Inhibitor 1 tablet of cOmplete^TM^ protease inhibitor cocktail (Roche) for each 10mL of buffer]. The proteins bound to the beads were boiled for 10 minutes in 1X Laemmli buffer containing 2-Mercaptoethanol and loaded in 10% SDS-Page electrophoresis until the sample reached the separation gel.

#### Protein Digestion and Sample Preparation

Protein digestion and mass spectrometry analyses were performed at the Proteomics Platform of the CHU de Québec Research Center (Quebec, Canada). Protein bands excised from gels were washed, reduced (10 mM DTT), and alkylated (55 mM iodoacetamide). Digestion was carried out with 126 nM trypsin (Promega) at 37 °C for 18 h. Peptides were extracted with 1% formic acid, 2% acetonitrile, followed by 1% formic acid, 50% acetonitrile. Extracts were pooled, dried by vacuum centrifugation, and resuspended in 10 µL of 0.1% formic acid.

#### Mass Spectrometry

Peptides were analyzed by nano-LC-MS/MS on a Dionex UltiMate 3000 system coupled to an Orbitrap Fusion mass spectrometer (Thermo Fisher Scientific). Samples were loaded onto a 5 mm × 300 µm C18 PepMap precolumn at 20 µL/min in loading solvent (2% acetonitrile, 0.05% TFA). Separation was performed on a 50 cm × 75 µm PepMap Acclaim column with a 5%–40% solvent B gradient (80% acetonitrile, 0.1% formic acid) over 200 min at 300 nL/min. Data were acquired in data-dependent mode (XCalibur 4.3) with full-scan MS (350–1800 m/z) at 120,000 resolution, AGC target of 4 × 10⁵, and maximum injection time of 50 ms. MS/MS fragmentation (HCD, 35% collision energy) was performed on the most intense ions (3 s cycle), with dynamic exclusion of 20 s (10 ppm tolerance). MGF files generated by Proteome Discoverer 2.3 were searched with Mascot 2.5.1 against a contaminant and *Leishmania braziliensis* Uniprot database (8153 entries). Parameters included trypsin digestion, fragment ion tolerance of 0.60 Da, and parent ion tolerance of 10 ppm. Fixed modification: carbamidomethylation (C). Variable modifications: deamidation (N, Q), oxidation (M), methylation/dimethylation (R). Protein and peptide identifications were validated using Scaffold 5.0.1. Peptides were accepted at >89% probability (FDR <1%), and proteins at >99% probability (FDR <1%) with ≥2 peptides (Protein Prophet algorithm). Significance was determined using One-Way ANOVA in Scaffold 5.0.1. (p < 0.05).

### Protein/RNA immunoprecipitation

Immunoprecipitations were conducted as described previously by Walrad (42,43) with modifications. Briefly 1×10^9^ log-phase promastigotes were harvested, washed twice with PBS and resuspended in 1mL of IP-Lysis Buffer (44). This material was immediately frozen in liquid nitrogen, defrosted and sonicated for 2 minutes twice (or until most of the parasites were lysed) and centrifuged 10,000 × *g* for 4 min at 4 °C. Supernatant was then incubated with 1mg of Pierce™ Anti-HA Magnetic Beads (Thermo Scientific™) for 2 hours at 4 °C under rotation. For the negative control (untagged parental cell line), magnetic beads were blocked with 100 μg of HA peptide for 30 minutes at 4 °C under rotation in IP-Lysis Buffer. Magnetic beads were washed at least 7 times to remove unbound proteins and divided in two tubes. Half of the material was destined to RNA extraction and DNAse treated as described in “RNA extraction section”. The other half was further divided in two tubes. One tube was incubated with 2x Laemnli buffer at 95°C for five minutes to extract the proteins from the beads. The other half was treated with 25 μg of DNAse and protease-free RNAse (Thermo Scientific™) in IP-Lysis Buffer without RNAse inhibitor by incubation at 22°C for 15 minutes. Magnetic beads were then washed three more times and incubated with 2x Laemnli buffer at 95°C for five minutes. The presence of the tagged protein in the eluate material was evaluated by immunoblotting using 1:5000 anti-HA antibody (Thermo Scientific™) in PBST-1% milk solution for 1 hour at 4°C, followed by incubation with 1:10,000 anti-Rabbit IgG (Promega) in PBST-1% milk solution overnight at 4°C. Membranes were incubated with Amersham ECL Prime Western Blotting Detection Reagents (Cytiva) and exposure and acquisition were conducted in BioRad ChemiDoc Imaging System.

### RNAseq

Total RNA, extracted as previously specified, was sent for processing by a specialized company (Macrogen, Inc.) that prepared the libraries (Illumina Ribo-Zero plus rRNA Depletion & TruSeq Stranded Total RNA Library Prep Gold Kit) and then proceeded to sequencing (Illumina platform), generating approximately 40 million short (101bp), paired and strand-specific reads per library. Mapping was performed using the bowtie2 program (version 2.3.2) (45), adding the parameters – local (excluding the need for end-to-end read alignment) and -N 1 (allowing a mismatch by alignment), using as reference the most recent version of the *Leishmania braziliensis* MHOM/BR/75/M2903 genome available in TriTrypDB (version 66) (46). The htseq-count program (part of the HTSeq package, version 0.12.4) (47), was used for counting, considering all reads (i.e., both reads that aligned unique and multiple times, parameter --nonunique=all) and the standard mode for assigning reads to features (parameter --mode=union). The R package DESeq2 (version 1.20.0) (48), was used for differential expression analysis, and genes with an adjusted p-value <0.05 were considered as differentially expressed. Additionally, a fold-change cutoff (FC > 1.5) was applied to prioritize genes with the greatest difference in expression between compared groups.

### Mass Spectrometry analysis of RIP samples

Sample provided in SDS-PAGE buffer was run approximately 1 cm into gel and stained with NBS SafeBLUE stain. In-gel digestion was performed with the addition of 0.2 μg sequencing-grade, modified porcine trypsin (Promega V5111), following reduction with dithioerythritol and alkylation with iodoacetamide. Digests were incubated overnight at 37°C. Peptides were extracted by washing three times with aqueous 50% (v:v) acetonitrile containing 0.1% (v:v) trifluoroacetic acid, before drying in a vacuum concentrator and reconstituting in aqueous 0.1% (v:v) trifluoroacetic acid.

Peptides were loaded onto EvoTip Pure tips for nanoUPLC using an EvoSep One system. A pre-set 100 SPD gradient was used with an 8 cm EvoSep C_18_ Performance column (8 cm x 150 μm x 1.5 μm). The nanoUPLC system was interfaced to a timsTOF HT mass spectrometer (Bruker) with a CaptiveSpray ionisation source. Positive PASEF-DIA, nanoESI-MS and MS^2^ spectra were acquired using Compass HyStar software (version 6.2, Bruker). Instrument source settings were: capillary voltage, 1,500 V; dry gas, 3 l/min; dry temperature; 180°C. Spectra were acquired between *m/z* 100-1,700. DIA windows were set to 25 Th width between *m/z* 400-1201 and a TIMS range of 1/K0 0.6-1.60 V.s/cm^2^. Collision energy was interpolated between 20 eV at 0.65 V.s/cm2 to 59 eV at 1.6 V.s/cm^2^.

DIA data were searched using DIA-NN (1.8.2.27) against the *Leishmania braziliensis* subset of Trytiyp-DB appended with common proteomic contaminants. Search criteria were set to maintain a false discovery rate (FDR) of 1% with heuristic protein inference. High-precision quant-UMS was used for extraction of quantitative values within DIA-NN. DIA-NN results were pivoted to protein-centric values using KINME (5.1.2), then filtered to protein identification q-values <0.01 and a minimum of two peptide matches per accepted protein group. Differential abundance testing was performed for pairwise comparisons using Limma via FragPipe-Analyst run in a local installation of R-shiny. Sample minimum imputation was applied without normalization. Local and tail area-based multiple test correction was used to generate adjusted p-values.

### GO analysis

The list of accession numbers was converted in Gene ID in Uniprot website. A list of IDs search was conducted in tritrypDB website, and the results were sent to gene ontology enrichment of biological processes analysis also in tritrypDB. A p-value of 0.05 was used as cutoff of significance. The retrieved terms, fold-change enrichment, number of genes in the term and p-value were used to generate a heat map using GraphPad Prism 10.

### Interactions networks

Interactions networks for the proteins identified in the pull-down assay for both *lncRNA45*WT (18 proteins) and *lncRNA45*MUT (13 proteins) were retrieved from the STRING database (32) using all available evidence types (i.e., Textmining, Experiments, Databases, Co-expression, Neighborhood, Gene Fusion and Co-occurrence). Cytoscape program (49) was used to highlight proteins and interactions shared between the networks, and an interactive version of both networks was submitted to the NDEx database (50).

## RESULTS

### Assembly of Trypanosomatid genomes for comparative analysis

In this study we selected eleven trypanosomatids genomes, pathogenic and nonpathogenic ones, for the search of RNA conserved secondary structures. We observed that for most of them, the percentage of Euglenozoa orthologues (complete sequence of the 130 orthologues) using BUSCO methodology was above 90%. This verifies the quality of genome assembly and annotation, with exception of the genomes of *Bodo saltans* and *Trypanosoma cruzi* (Table 1). Despite their lower conservation levels, the genomes of *Bodo saltans* and *Trypanosoma cruzi* were included in the analysis to ensure representation of diverse evolutionary lineages, as their complete orthologue percentages remained higher than fragmented or missing sequences. The percentage of complete orthologues (C(%), Table 1) exceeded the percentage of fragmented (F(%), Table 1) or missing sequences (M(%), Table 1).

**Table 1-.** BUSCO analysis of trypanosomatids’ genomes. The universal orthologues used for this analysis were obtained from the BUSCO euglenozoan Odb10 dataset.

| Species<sup>a</sup> | BUSCO (euglenozoa – 130)<sup>b</sup> |  |  |  |
| --- | --- | --- | --- | --- |
|  | C(%) | CD(%) | F(%) | M(%) |
| *Leishmania major* Friedlin | 98.5 | 0 | 0 | 1.5 |
| *Leishmania donovani* BPK282A1 | 97.7 | 0 | 0 | 2.3 |
| *Leishmania infantum* JPCM5 | 97.7 | 0 | 0 | 2.3 |
| *Leishmania amazonensis* MHOM/BR/1973/M2269 | 96.9 | 0 | 0.8 | 2.3 |
| *Leishmania panamensis* MHOM/COL/81/L13 | 96.9 | 0 | 1.5 | 1.6 |
| *Crithidia fasciculata* CF-C1 | 96.2 | 0 | 2.3 | 1.5 |
| *Leishmania braziliensis* MHOM/BR/75/2903 | 93.8 | 0 | 4.6 | 1.6 |
| *Trypanosoma brucei* TREU927 | 93.1 | 7.7 | 3.1 | 3.8 |
| *Bodo saltans* LakeKonstanz | 72.3 | 0.8 | 9.2 | 18.5 |
| *Trypanosoma cruzi* CL Brener Non-Esmeraldo-like | 66.9 | 0 | 3.8 | 29.3 |
| *Trypanosoma cruzi* CL Brener Esmeraldo-like | 54.6 | 0 | 3.1 | 42.3 |

<sup>a</sup>Species of Trypanosomatids (genome files from TriTrypDB v44)
<sup>b</sup>BUSCO analysis results: (Complete - C(%)), percentage of 130 Euglenozoa universal single copy orthologues identified as full sequence orthologues in the listed genomes. (Complete Duplicated - CD(%)) full sequence found in more than one position in the genome. (Fragmented – F(%)) only partial sequence was found or; (Missing – M(%))sequence not found.

We aligned these 11 trypanosomatid genomes against the *Leishmania braziliensis* MHOM/BR75/M2903 genome using Lastz (26), and the pairwise alignment was combined into a *L. braziliensis* based multiple alignment using MultiZ (51) and the UCSC tool chain (see Methods for details). Details of genome quality, including percentages of complete, fragmented, and missing orthologues, are presented in Table 1.

### Prediction of conserved structures

To predict conserved RNA structures in the *L. braziliensis* genome we used MultiZ/roast to form a multiple alignment of 10 genomes of *Leishmania* species, covering 99.3 % of the *L. braziliensis* genome. From the multiple alignment we constructed 12,969 alignment windows that overlap with the lncRNA annotation. On these windows we predicted conserved RNA structures with RNAz (52). The screen with RNAz yields 631 ncRNA candidates with a support vector machine (SVM) score >=0, which is the threshold used by RNAz to classify the window as a conserved RNA structure.

Infernal (version 1.1.2) (53) / Rfam (version 14.1) (54) predicted 177 known structured RNAs in the *L. braziliensis* genome and to substantiate our findings we overlapped and compared the 177 known structures with the RNAz predictions. 226 of the 12,969 alignment windows overlap 64 of the known structures with at least 40 bases. We analyzed the 12,969 alignment windows in several ways and determined that the average pairwise sequence identity and the number of organisms/sequences in each alignment window were the most efficient filters (Figures S1 and S2). We thus filtered the alignment windows by average pairwise sequence id >= 70% and 6>= organisms in the alignment (if 7 or more sequences are found to be valid in the input alignment, RNAzWindow will automatically select the 6 best sequences based on optimal pairwise sequence identity). After this filter, we are left with 6,035 alignment windows, of which 206 overlap known RNA structures.

To estimate the false discovery rate for the RNAz predictions, we shuffled the 6,035 alignment windows. Each window was shuffled 100 times under conservation of di-nucleotide contents and the SVM scores of the shuffled windows were used as background to yield a p-value for the RNAz predictions (Supplementary Figure 3). Based on the p-values of the 206 windows overlapping the known ncRNAs we chose a cut-off for the SVM score of 0.5, which corresponds to p-value <= 0.004 (Supplementary Figure 4 and Supplementary Figure 3). To avoid GC-bias of the individual candidate, we used the 100 shuffled windows of each candidate as its own background and required that all 100 shuffled windows have an SVM score lower than the candidate’s score. This ensures that the per-candidate FDR is < 0.01. The filtering based on the background p-value <= 0.004 and per candidate FDR < 0.01, leads to a final set of 142 candidates, of which 104 are novel and 38 are overlapping known ncRNAs, conversely, we recover 19 of 54 expressed, highly conserved ncRNAs (38 of 206 windows, see Figure 1).

**Figure 1-.**
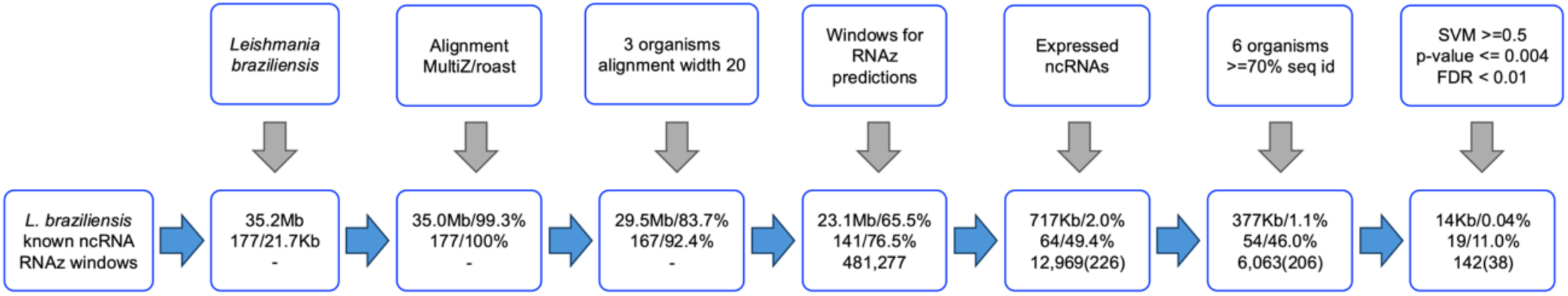
The filtering process of the multiple alignment covering 99.3% of the genome down to 12,969 multiple alignment windows on which we predict 142 structured RNA candidates where 104 are novel while 38 overlap the 177 known structured RNAs predicted by Infernal (version 1.1.2) (25) / Rfam (version 14.1) (26) predicts 177 known. After each step, we show statistics in the bottom panels. The three rows in the bottom panels are coverage of the Lbr genome in bases and percentage, coverage of the 64 known and expressed ncRNAs, and the number of structure predictions.

In the final selection, we focused on candidates located between two CDSs on the same strand, which is true for a good portion of the known ncRNAs (∼55% within 10,000 and ∼38% within 5,000 bases). We found that 74 of our 104 novel candidates are located between two CDSs (<=5000). The ncRNAs that are preferentially expressed in the amastigote stage, were chosen to be functionally analyzed. This choice was based on the possibility that these ncRNA play a role in the intracellular survival of the parasite in the disease-relevant stage. Thus, the investigation may provide novel insights into the mechanisms orchestrating parasite-host interactions.

### Predicted point mutations that can affect the structures of target ncRNAs

Considering its coverage, size, genomic position, and differential expression, the lncRNA *LbrM2906_26_lncRNA45* (*lncRNA45*) was identified as a candidate for functional characterization. This lncRNA harbors a conserved structured region, referred to as Locus 709.

After target selection, *in silico* prediction of mutations that could affect the RNA secondary structure, as well as its stability or interactions with other macromolecules, was performed using the *RNAsnp* tool. The nucleotide substitution of a cytosine by a guanine at position 50 of *lncRNA45*, in the conserved Locus709, was predicted in silico, to potentially change the conformation of this lncRNA secondary structure which could impair its function. *RNAsnp* reported several possible mutations that would alter the wild-type structure. Among these, the mutation of C to G at position 50 had the lowest p-value of 0.004. Interestingly, many of the other SNPs induced a structure quite different for the wild-type structure, for example G to A at position 76 (Figure 2). It is possible that the functional structure is the one obtained from G76A, and that we destabilize this structure by the induced mutation of C50G; however, this would require further investigation beyond the scope of the current study.

**Figure 2:**
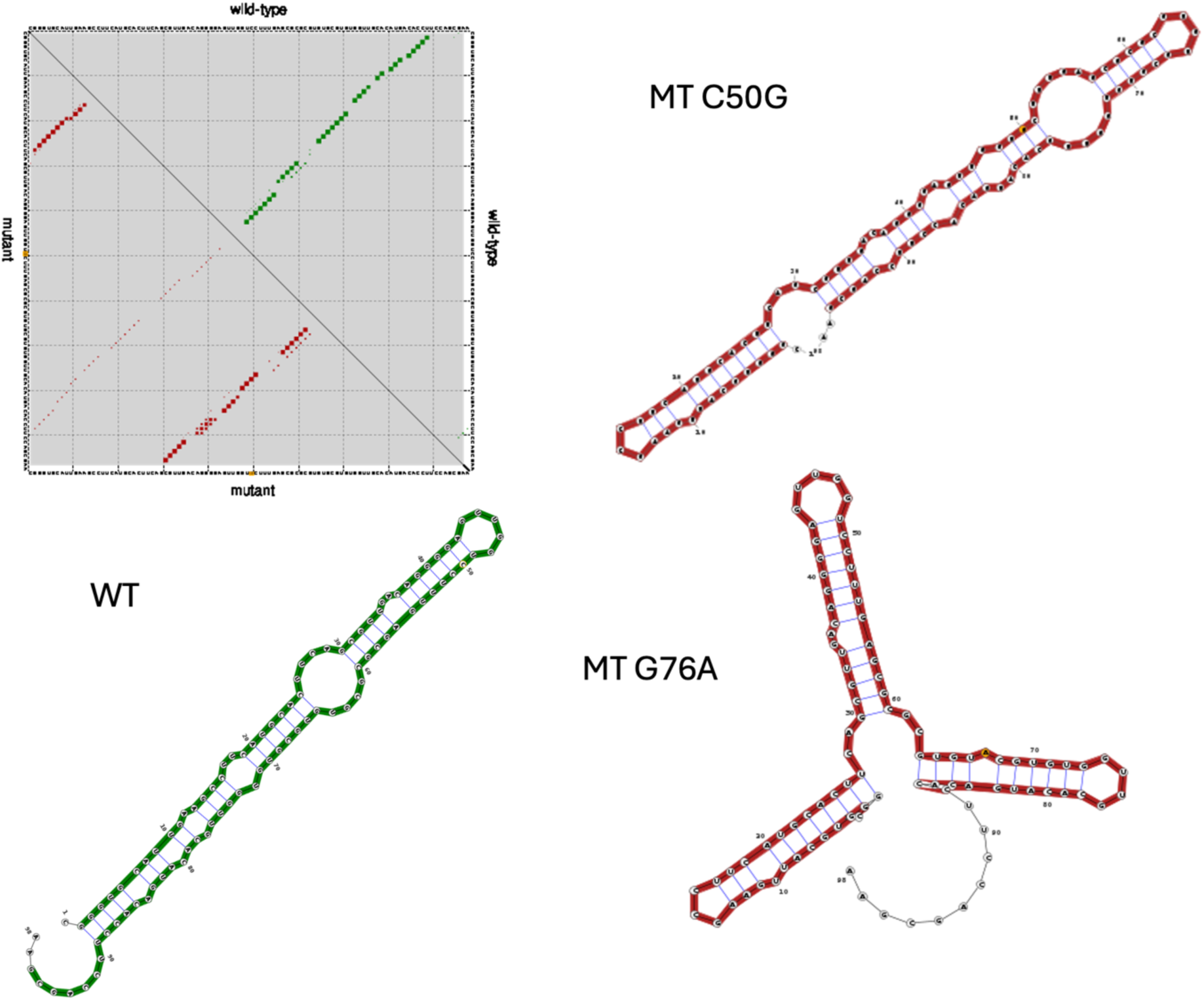
Outcome of running RNAsnp on the tested candidate. Top left, base pair probabilities of wildtype (WT) and mutant (MT) after the mutation of C to G at position 50. Top right, MFE structure of the folded MT RNA. Bottom left, MFE structure of the WT. Bottom right, MFE structure of the folded RNA after mutating G to A at position 76.

### *lncRNA45* is processed and localizes in the cytoplasm of *L. braziliensis*

In a previous publication by our group (17), *lncRNA45* was identified in the *L. braziliensis* M2903 transcriptome and predicted to be a non-coding RNA (ncRNA) by at least two predictors: ptRNApred (classified as RNaseP family) and RNAcon (classified as Intron-gp-1). The predicted size of *lncRNA45* is 408 nt, and it was found to be located in an intergenic region of chromosome 26 (coordinates: 296,265 - 296,672) on the minus strand, which is the same strand sense as the PTU it belongs to (Figure 3A) (17).

**Figure 3-.**
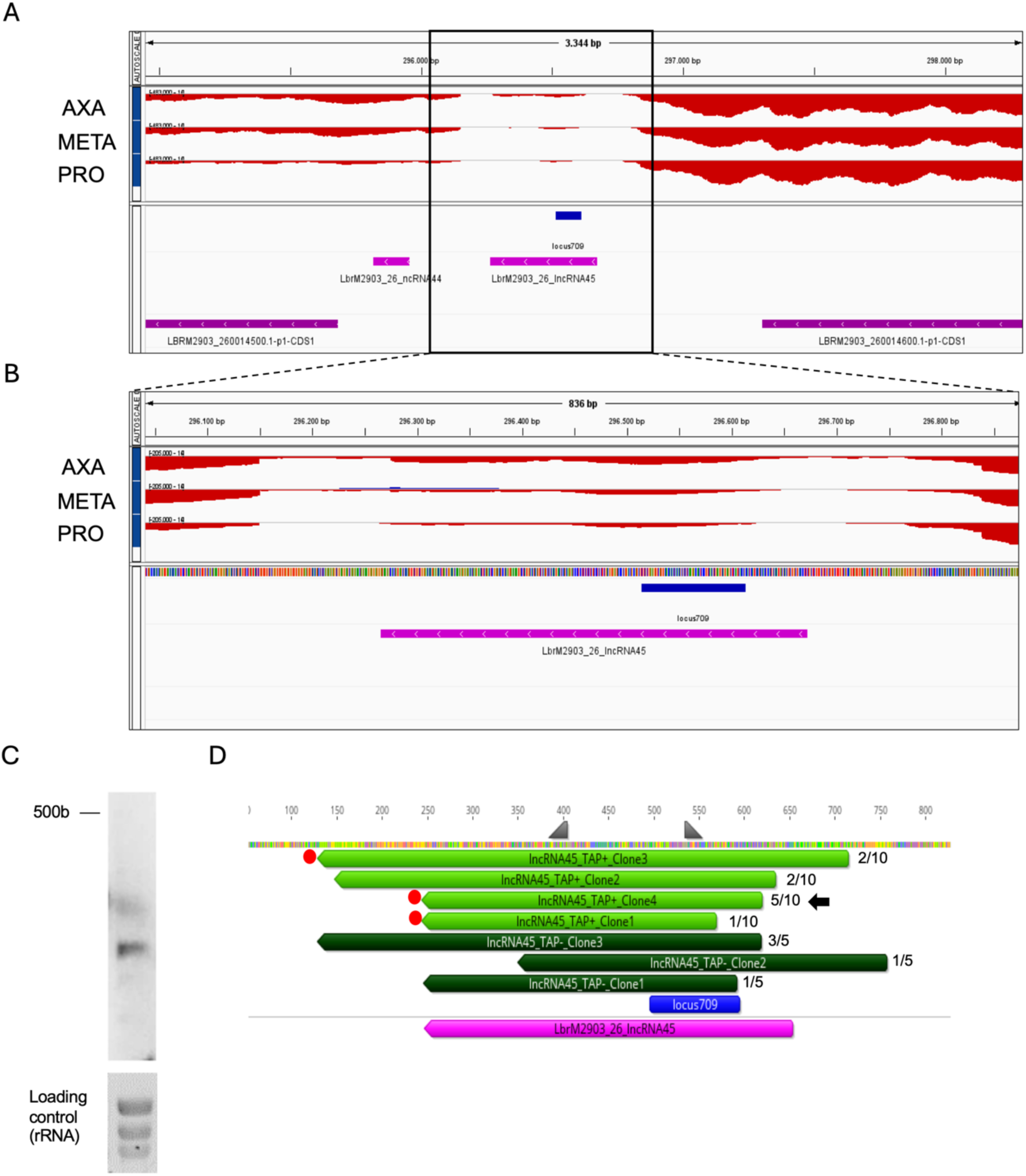
Characterization of LbrM2903_26_lncRNA45 (lncRNA45). (A) IGV representation of genomic regions on chromosome 26 of *L. braziliensis* MHOMBR75M2903, showing lncRNA44 and lncRNA45 (magenta), two coding sequences (CDSs) - hypothetical protein LBRM2903_260014500.1 and NMD3 family protein LBRM2903_260014600.1 (purple), and the predicted locus 709 (blue). (B) RNA-seq coverage (red stacking bars) of lncRNA45 (magenta) in axenic amastigotes (AXA), metacyclic promastigotes (META), and procyclic promastigotes (PRO). The target locus 709 is indicated in blue. (C) Detection of lncRNA45 as a transcript smaller than 500 bp in total RNA extract from *L. braziliensis* M2903 procyclic promastigotes, by northern blotting using a radiolabeled specific probe. rRNA served as a loading control. (D) Characterization of lncRNA45 size and processing pathways using RNA circularization. Total RNA from *L. braziliensis* M2903 promastigotes was extracted and treated with tobacco acidic phosphatase for decapping (TAP+) or left untreated (TAP-), followed by circularization with T4 RNA ligase. Complementary circular DNA (ccDNA) was synthesized by reverse transcription using a primer binding 200 nt from the 5’ end of lncRNA45. PCR amplification with primers lncRNA45_cF1 and lncRNA45_cR1 (shown in gray) was performed using ccDNA as a template, and PCR products were cloned into pGEM-T for sequencing (Hang, Deng et al., 2015). Sequences from three cDNA clones derived from TAP- (dark green) and four from TAP+ (light green) were analyzed. Poly(A) tails (red dots) were detected in 3 out of 7 transcript profiles, all derived from TAP+ templates. The most representative transcript profile among the clones is indicated by a black arrow.

*LncRNA45* is situated between two coding sequences (CDSs): an upstream gene encoding a putative NMD3 family protein (LbrM2903_260014600.1), and a downstream gene coding for a hypothetical protein (LbrM2903_260014500.1) (55) (Figure 3A). The RNA coverage profile strongly suggests that *lncRNA45* is not part of the untranslated region (UTR) of either of these CDSs (Figure 3B). In addition, the RNAseq coverage profile indicates that *lncRNA45* is preferentially expressed in amastigotes (Figure 3A and 3B), showing a 1.53-fold higher abundance in amastigotes compared to other morphological forms. However, the RT-qPCR analysis did not confirm the RNA-seq results, revealing instead that *lncRNA45* is more abundant in promastigotes (PRO) than in other morphologies (Supplementary Figure 5C).

The existence of *lncRNA45* as an independent transcript in *L. braziliensis* M2903 total RNA extract was confirmed by northern blotting using sequence-specific radiolabeled probes (Figure 3C). A specific, unique band smaller than 500 bp was observed for *lncRNA45*, confirming the presence of this ncRNA transcript in *L. braziliensis*, consistent with the expected size of 408 nucleotides.

To confirm the size of *lncRNA45* and investigate its processing features following polycistronic transcription, an RNA circularization-based protocol (34) was employed (Supplementary Figure 5A). Total RNA from *L. braziliensis* M2903 was decapped (TAP+) or not (TAP-) using tobacco acidic phosphatase (TAP), circularized, and reverse transcribed using target-specific primers. Primers directed towards the 5’ and 3’ ends were used to amplify a product containing the transcript ends and its modifications (poly(A) tail and spliced leader, if present). A distinct and specific band was observed in both TAP+ and TAP-reactions (Supplementary Figure 5B). These bands were cloned into pGEM-T and sequenced. Fifteen clones (ten from TAP+ and five from TAP-reactions) were collected, amplified with M13-F and M13-R primers and sequenced. Nine clones presenting different fragment sizes after amplification were sequenced and mapped to the locus of *lncRNA45* on *L. braziliensis* M2903 chromosome 26.

*In silico* predictions suggested that *lncRNA45* is 407 nucleotides long, but the circularization assay revealed 7 different transcript profiles ranging from 325 to 585 bp (Figure 3D). Among these, 4 cDNA clones lacked a poly(A) tail in the 3’ UTR, while 3 clones had a poly(A) tail ranging from 14 to 17 adenines (Figure 3D). No spliced leader sequence was observed in the 5’ UTR of *lncRNA45*, suggesting that it may be capped by a mechanism other than spliced leader-mediated capping. Interestingly, three different transcript sizes lacking polyadenylation were identified in the TAP-reactions, whereas three out of four clones sequenced from the TAP+ reactions were polyadenylated. The most abundant transcript in TAP+ samples (5 out of 10 clones) was 376 nucleotides long and had a 13-nucleotide poly(A) tail (indicated by the black arrow in Figure 3D), corresponding to the region of highest coverage observed for *lncRNA45* (Figure 3D). Notably, except for *lncRNA45*_TAP+_Clone1, all other identified *lncRNA45* isoforms contained the conserved structured Locus 709 identified in comparative analysis.

The localization of *lncRNA45* was determined using RNA fluorescence *in situ* hybridization (FISH) with specific probes binding along its entire length. A clear cytoplasmic localization was observed, with no signal co-localizing with *L. braziliensis* nuclei in any analyzed cells (Figures 4A-E). In some cells, multiple dots were observed, but due to experimental limitations, it was not possible to confirm whether each dot represents one or more *lncRNA45* molecules. Interestingly, *lncRNA45* was detected in both parental and daughter cells, even during cell division.

**Figure 4-.**
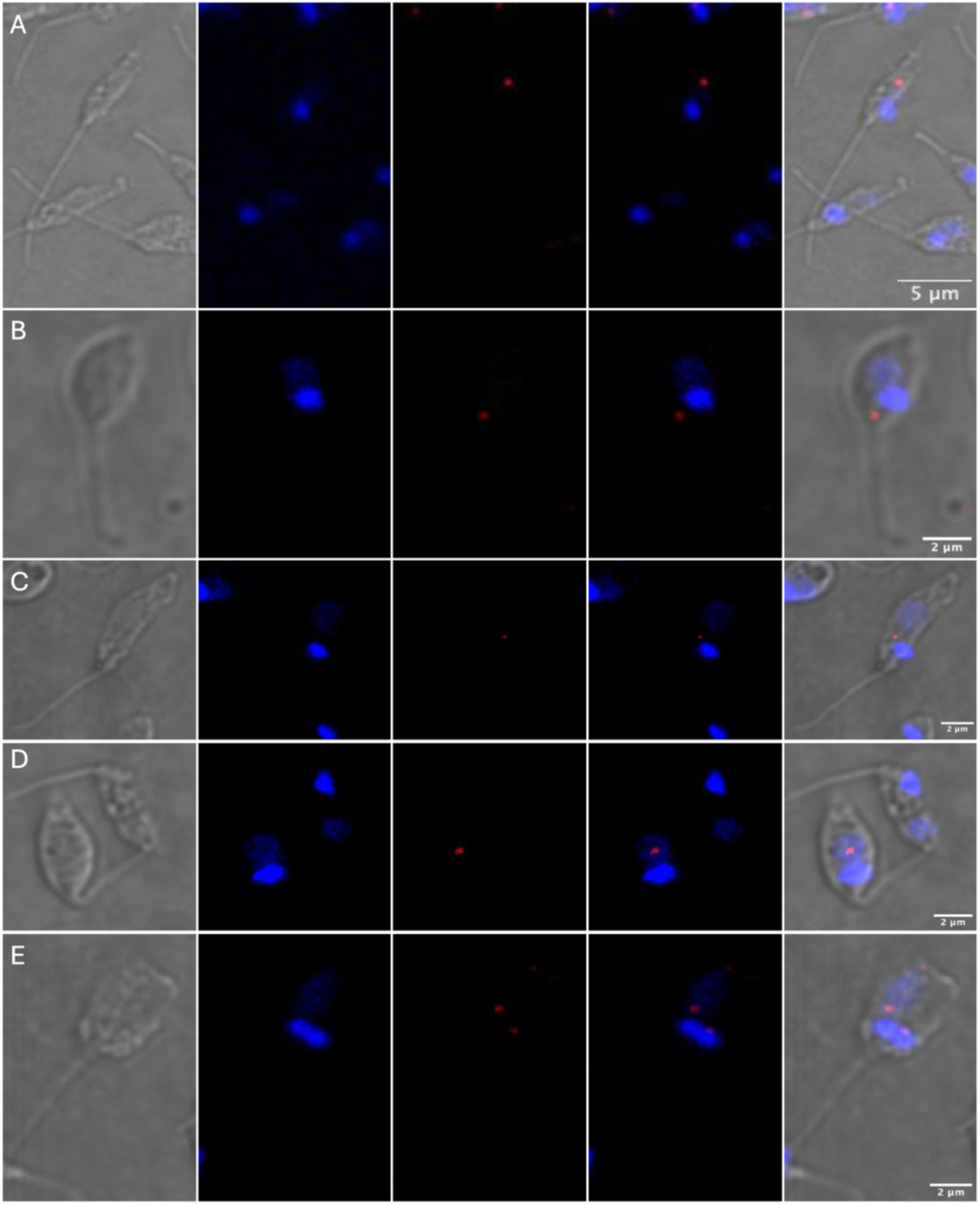
LncRNA45 localizes to the cytoplasm of *L. braziliensis*. RNA FISH was conducted using specific probes spanning the entire length of lncRNA45. Following probe incubation, permeabilized cells were treated sequentially with pre-amplifier and amplifier solutions for fluorophore binding. Images were captured using a Zeiss multiphoton microscope at 100x magnification. The results showed that lncRNA45 (red dots) exclusively localizes to the cytoplasm of *L. braziliensis*, with no co-localization observed with nuclei (stained with DAPI, shown in blue) in any of the cells.

### *lncRNA45* knockout results in reduced parasite density in culture

To investigate the biological function of *lncRNA45*, *L. braziliensis* M2903 cell lines knocked out (Δ*lncRNA45*) for *lncRNA45*, or overexpressing (OE_*lncRNA45*) this transcript, were generated. Using CRISPR/Cas9, *lncRNA45* was deleted from the *L. braziliensis* M2903 genome (30,31,38). Null mutants were successfully obtained, and the complete loss of *lncRNA45* was confirmed by PCR (Supplementary Figure 6A and Supplementary Figure 6E – PCR A).

*L. braziliensis* M2903 overexpressing *lncRNA45* (OE_*lncRNA45*) was obtained by transfection with the pSSU-*lncRNA45*-Sat plasmid and selection with nourseothricin (Supplementary Figure 6B and Supplementary Figure 6F – PCR B). As control, *L. braziliensis* M2903 was also transfected with pSSU-GFP-Sat plasmid (OE_mock) (Supplementary Figure 6C and Supplementary Figure 6F – PCR C).

To investigate whether parasite fitness was impaired, these transfectants (Δ*lncRNA45*, OE_*lncRNA45*, and OE_mock) along with the parental cell line, were subjected to assays mimicking key steps of the *Leishmania* life cycle. Promastigote growth was measured by adjusting the culture to 2 × 10^5^ promastigotes/mL and counting for at least 7 days or until the culture reached the stationary phase. Area under the curve analysis showed significantly reduced growth for Δ*lncRNA45*, with a cell density of 3.6 × 10^7^ promastigotes/mL on the 7th day of culture, compared to 5.5 × 10^7^ promastigotes/mL for the parental cell line (Figure 5A). No significant differences were observed for OE_*lncRNA45* or OE_mock, compared to the parental cell line (One-way ANOVA and Tukey’s multiple-comparison test, p < 0.05).

**Figure 5-.**
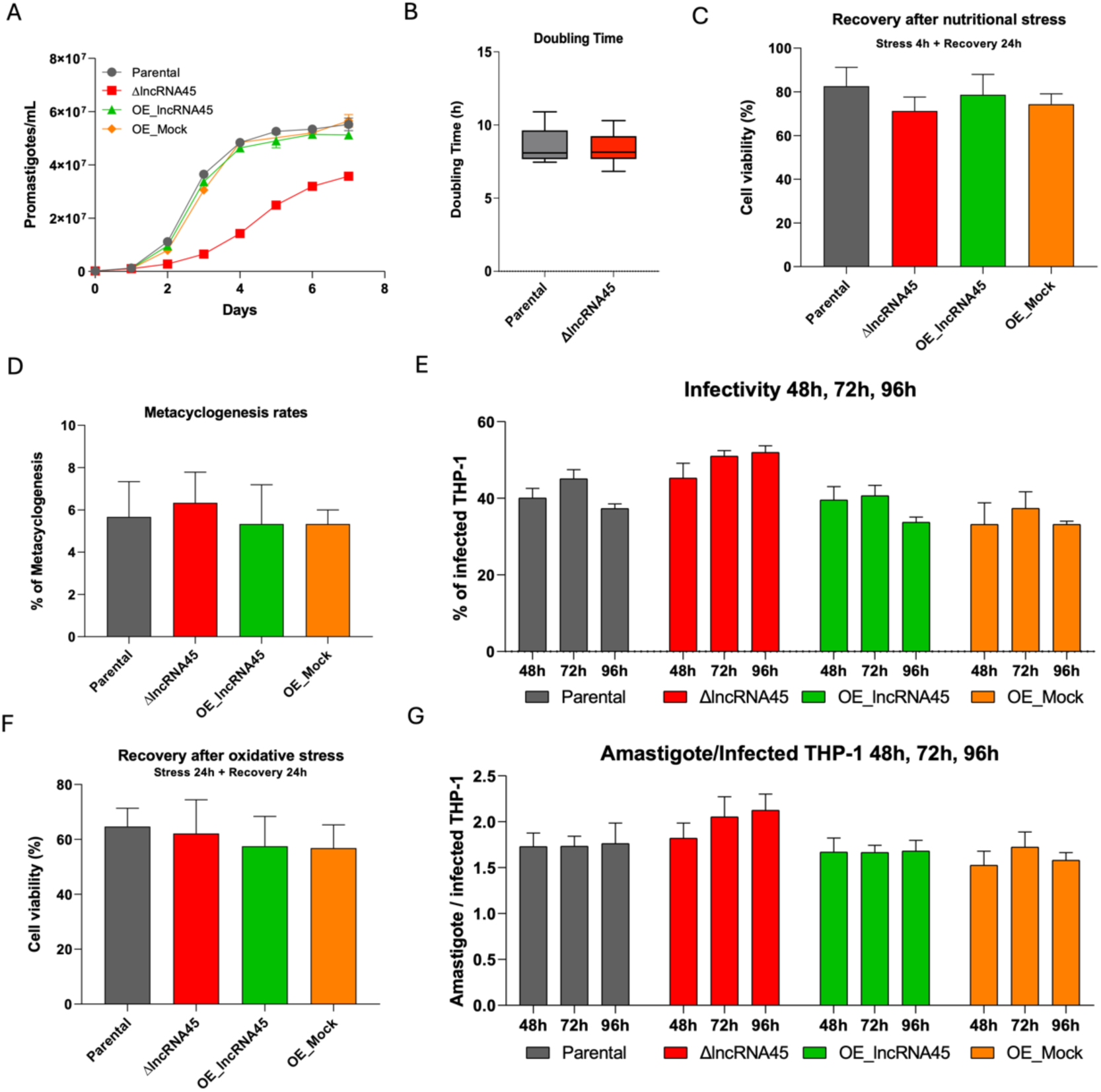
Phenotypic characterization of *L. braziliensis* M2903 ΔlncRNA45 (red), OE_lncRNA45 (green), and OE_Mock (orange) transfectants compared to the parental cell line (gray). (A) Promastigote growth for each cell line was determined by counting the culture for seven consecutive days. A significant reduction in growth was observed in ΔlncRNA45 by area under the curve (AUC) analysis compared to the other cell lines (One-way ANOVA and Tukey’s multiple-comparison test, p < 0.05). (B) Doubling time was determined for ΔlncRNA45 in comparison to the parental cell line in four biological replicates (Non-parametric T-test, p < 0.05). (C) The capacity to recover from nutritional stress induced by incubation with PBS for 4 hours was measured by MTT assay for the transfectants relative to non-stressed cells and the parental line. No significant differences were observed (One-way ANOVA and Tukey’s multiple-comparison test, p < 0.05). (D) The percentage of metacyclic promastigotes in late stationary culture of transfectants compared to the parental cell line was determined by Ficoll purification, and no significant differences were observed (One-way ANOVA and Tukey’s multiple-comparison test, p < 0.05). (E) The capacity of transfectants to recover from oxidative stress induced by incubation with H_2_O_2_ for 24 hours was measured by MTT assay relative to non-stressed cells and compared to the parental line. No significant differences were observed (One-way ANOVA and Tukey’s multiple-comparison test, p < 0.05). (F) The infectivity of the transfectants to THP-1 macrophages and the multiplying capacity as intracellular amastigotes was evaluated in comparison to the parental cell line. No significant differences in the percentage of infected cells (G) and the number of amastigotes per infected macrophage (H) were observed at any time point (One-way ANOVA and Tukey’s multiple-comparison test, p < 0.05).

To determine if the reduced cell density observed after 7 days for the Δ*lncRNA45* cell line was due to reduced replication capacity, doubling time was measured by adjusting the culture to 1 × 10^6^ promastigotes/mL and counting after 24 hours. This procedure was repeated for 4 days, and no significant differences in doubling time were observed for Δ*lncRNA45* compared to the parental cell line (Figure 5B) (Non-parametric t-test, p < 0.05).

When inside the vector, procyclic promastigotes migrate to the anterior portion of the phlebotomine gut and face an environment with reduced nutrient availability. The capacity to resist nutritional stress was evaluated for Δ*lncRNA45* and OE_*lncRNA45* by incubating these cells in PBS for 4 hours, followed by incubation in M199 for 24 hours. The percentage of viable cells after stress was determined relative to non-stressed cells and compared to the parental line. No significant differences were observed in the capacity to recover from nutritional stress for any of the transfectant lines compared to the parental cell line (Figure 5C) (One-way ANOVA and Tukey’s multiple-comparison test, p < 0.05).

To infect the mammalian host, procyclic promastigotes need to differentiate into infective metacyclic promastigotes, which are regurgitated at the lesion site during the blood meal and infect mammalian host cells. Using a Ficoll gradient, we purified metacyclic promastigotes from late stationary phase cultures and determined the percentage of metacyclic promastigotes compared to procyclic promastigotes for Δ*lncRNA45*, OE_*lncRNA45*, as well as the parental and OE_mock cell lines. The percentages of metacyclic promastigotes in culture (% of metacyclogenesis) observed for the parental, Δ*lncRNA45*, OE_*lncRNA45*, and OE_mock cell lines were 5.7% ± 1.7 (mean ± SEM), 6.3% ± 1.5, 5.3% ± 1.9, and 5.3% ± 0.6, respectively, with no significant differences observed between the cell lines (Figure 5D) (One-way ANOVA and Tukey’s multiple-comparison test, p < 0.05).

The capacity to infect THP-1-derived macrophages and the replication of intracellular amastigotes were assessed for Δ*lncRNA45*, OE_*lncRNA45*, and OE_mock in comparison to the parental cell line. No significant differences in the percentage of infected macrophages (Figure 5E) or in the number of amastigotes per macrophage (Figure 5G) were observed up to 96 hours post-infection for any transfectant compared to the parental cell line (One-way ANOVA and Tukey’s multiple-comparison test, p < 0.05).

Once inside the macrophages, *Leishmania* parasites are exposed to reactive oxygen species, which they must survive to replicate and establish infection. The capacity to recover from oxidative stress caused by hydrogen peroxide (H_2_O_2_) was evaluated for Δ*lncRNA45*, OE_*lncRNA45*, and OE_mock in comparison to the parental cell line. After incubation with H_2_O_2_ for 24 hours, parasites were allowed to recover in M199 media for another 24 hours. The percentage of recovery was determined in comparison to non-stressed parasites, and no significant differences were observed for the transfectant lines compared to the parental cell line (Figure 5F).

### *lncRNA45*-protein interaction is modified in the presence of C50G substitution in Locus 709

The observation that the absence of *lncRNA45* affects *L. braziliensis* promastigote fitness, along with the fact that this ncRNA is differentially expressed between morphologies and has a conserved structured locus, strengthens the hypothesis that this lncRNA may be involved in gene expression regulation in this parasite. To investigate which pathways *lncRNA45* might regulate, an *in vitro* pulldown assay was conducted to identify the proteins that interact with this lncRNA.

To determine if the RNA-protein interaction is dependent on the secondary structure formed by Locus 709, the pulldown was performed using both the wild-type sequence of *lncRNA45* (*lncRNA45*WT) and a sequence containing the C50G substitution (*lncRNA45*MUT). The *lncRNA45*WT and *lncRNA45*MUT sequences were fused to a 4xS1m aptamer by restriction enzyme cloning into the pUC57-4xS1m plasmid and sequenced to ensure that C50G was the only divergence between the sequences. Since S1m has an affinity for streptavidin-coated beads (SA-beads), *lncRNA45* was immobilized and exposed to log-phase *L. braziliensis* M2903 promastigote protein lysate. As a control, an S1m sequence without the *lncRNA45* fusion was used.

Proteins recovered after filter selection and significantly enriched in all three replicates of *lncRNA45*WT and *lncRNA45*MUT compared to the empty control filter were considered binding partners of these sequences. A total of 18 and 13 proteins were identified as binders for *lncRNA45*WT and *lncRNA45*MUT, respectively (Table 2, Figure 6A). Interestingly, the 13 proteins recovered from the *lncRNA45*MUT pulldown were common to the *lncRNA45*WT pulldown (Table 2). The other five proteins were lost in the presence of the C50G substitution in the Locus 709 sequence of *lncRNA45* (*lncRNA45*MUT) (Table 2, Figure 6A).

**Figure 6-.**
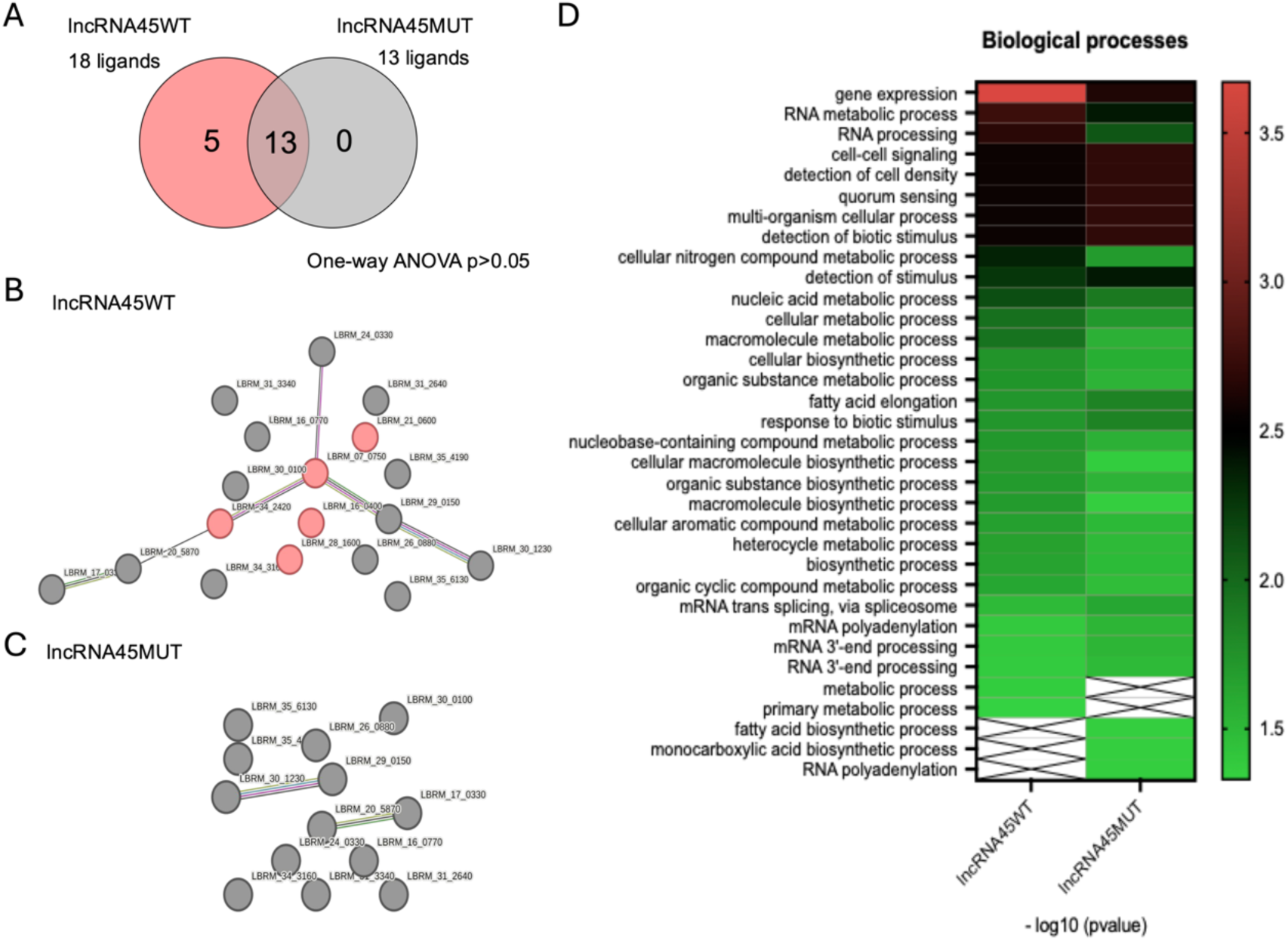
*In Vitro Pulldown of lncRNA45WT and lncRNA45MUT sequences*. (A) The lists of proteins that bind to lncRNA45WT (18 proteins) and lncRNA45MUT (13 proteins) were compared to identify common and specific ligands. In the presence of the C50G substitution in locus 709, five lncRNA45-protein interactions were lost. Thirteen proteins interacted with lncRNA45 in both the presence and absence of the C50G mutation. The protein-protein interaction network was determined for the proteins binding to lncRNA45WT (B) and lncRNA45MUT (C) using String. The interactive networks can be accessed for lncRNA45WT (https://www.ndexbio.org/#/network/0175a6c3-274c-11ef-9621-005056ae23aa?accesskey=e4fe291b38d4e92ee1b1585d11e0ca5bf624fdf6e669a3155c711dde39d39089) and for lncRNA45MUT (https://www.ndexbio.org/#/network/d3df8921-274a-11ef-9621-005056ae23aa?accesskey=e505803df77aa75dbf287c55f80c2226fb71bfc82ac437ab93cd47d4975ec1db). (D) Heat map of gene ontology (GO) enrichment analysis of proteins binding to lncRNA45WT and lncRNA45MUT was performed. The list of accession numbers retrieved from mass spectrometry was converted into *L. braziliensis* M2904 gene IDs using UniProt and then submitted to GO enrichment analysis of biological processes in TriTrypDB. A p-value cutoff of 0.05 was used. The resulting lists of enriched terms were classified based on significance (p-value).

**Table 2-.** lncRNA45-protein interactions observed in S1m in vitro pull-down.

| lncRNA45 |  |  |  |  |  |  |
| --- | --- | --- | --- | --- | --- | --- |
| Code | Description<sup>a</sup> | Accession Number<sup>b</sup> | GeneID M2904<sup>c</sup> | GeneID M2903<sup>c</sup> | Size<sup>d</sup> | pValue<sup>e</sup> |
| Lb1 | 40S ribosomal protein S9, putative | A4H519 | LbrM.07.0750 | LBRM2903_070014500 | 22 kDa | 95% (0.0029) |
| Lb2 | XRN 5'-3' exonuclease N-terminus, putative | A4H8G4 | LbrM.16.0400 | LBRM2903_160009500 | 108 kDa | 95% (0.011) |
| Lb3 | Lupus La protein homolog, putative | A4HBQ1 | LbrM.21.0600 | LBRM2903_210010600 | 37 kDa | 95% (0.029) |
| Lb4 | hypothetical protein, conserved | A4HGJ6 | LbrM.28.1600 | LBRM2903_280022100 | 80 kDa | 95% (0.014) |
| Lb5 | Nop53 (60S ribosomal biogenesis), putative | A4HMS1 | LbrM.34.2420 | LBRM2903_340031800 | 38 kDa | 95% (0.0080) |
| Lb6 | RNA-binding protein, putative | A4HI65 | LbrM.30.1230 | LBRM2903_300017800 | 67 kDa | 95% (0.014) |
| Lb7 | tRNA (Uracil-5-)-methyltransferase, putative | E9AIL9 | LbrM.20.5870 | LBRM2903_200073900 | 66 kDa | 95% (0.047) |
| Lb8 | DNA-directed RNA polymerase II subunit 3, putative | A4HH22 | LbrM.29.0150 | LBRM2903_290006800 | 37 kDa | 95% (0.037) |
| Lb9 | dihydrouridine synthase (Dus), putative | A4H8X8 | LbrM.17.0330 | LBRM2903_170009100 | 66 kDa | 95% (0.037) |
| Lb10 | Mitochondrial small ribosomal subunit Rsm22, putative | A4HD89 | LbrM.24.0330 | LBRM2903_240008300 | 110 kDa | 95% (0.031) |
| Lb11 | ISWI complex protein | A4HEU2 | LbrM.26.0880 | LBRM2903_260013600 | 80 kDa | 95% (0.026) |
| Lb12 | kinetoplast-associated protein, putative | A4HQA1 | LbrM.35.6130 | LBRM2903_350072700 | 14 kDa | 95% (0.041) |
| Lb13 | acetyl-CoA<br>carboxylase | A4HJT6 | LbrM.31.3340 | LBRM2903_310043300 | 241 kDa | 95%<br>(0.0053) |
| Lb14 | N-terminal region of<br>Chorein, a TM<br>vesicle-mediated<br>sorter, putative | A4H8K0 | LbrM.16.0770 | LBRM2903_160014400 | 604 kDa | 95%<br>(0.049) |
| Lb15 | Symplekin tight<br>junction protein C<br>terminal, putative | A4HJL9 | LbrM.31.2640 | LBRM2903_310035900 | 165 kDa | 95%<br>(0.0028) |
| Lb16 | replication factor C,<br>subunit 3, putative | A4HMZ2 | LbrM.34.3160 | LBRM2903_340040300 | 40 kDa | 95%<br>(0.011) |
| Lb17 | hypothetical protein,<br>conserved | A4HHV4 | LbrM.30.0100 | LBRM2903_300006000 | 24 kDa | 95%<br>(0.042) |
| Lb18 | hypothetical protein,<br>conserved | A4HPR3 | LbrM.35.4190 | LBRM2903_350052100 | 51 kDa | 95%<br>(0.0021) |

<sup>a</sup>Protein description based on TritypDB annotation.
<sup>b</sup>Uniprot accession number
<sup>c</sup>Gene identification in TritypDB
<sup>d</sup>Protein size in kilodalton (kDa)
<sup>e</sup>pValue and % of confidence obtained in the statistical analysis. One-Way ANOVA was used to compare S1m results of the negative control, the lncRNA45WT and lncRNA45MUT pull-downs. Unpaired T-test was used to compare S1m results of the negative control with the lncRNA52 pull-down. The differences were considered significant when $p < 0.05$ .
**Bold** labelling refers to proteins specifically binding to lncRNA45WT

Protein interaction network analysis was conducted on the protein interactors of *lncRNA45*WT (Figure 6B) and *lncRNA45*MUT (Figure 6C). A network composed of 6 proteins was observed when binding partners of *lncRNA45*WT were imputed (Figure 6B). When binding partners of *lncRNA45*MUT were imputed, however, this network was lost (Figure 6C). Indeed, two proteins (LbrM.07.0750 and LbrM.34.2420) of the five found to bind only the native sequence of *lncRNA45* are components of this network (Figure 6A).

To identify the pathways *lncRNA45* might be involved in, the list of binding partners for *lncRNA45*WT and *lncRNA45*MUT sequences was submitted to gene ontology (GO) enrichment analysis (Figure 6D). Twelve and seventeen terms were retrieved for *lncRNA45*WT and *lncRNA45*MUT, respectively. Eleven of these terms were shared between both protein lists. RNA processing (GO:0006396) was exclusively enriched in *lncRNA45*WT, while six terms were exclusively enriched in *lncRNA45*MUT (Figure 6D).

Since genome annotation can significantly impact GO analysis (56), we identified the orthologs of the *L. braziliensis* genes retrieved in the pulldown analysis within the *L. major* Friedlin genome, which has a more complete annotation to date, to compare the outcomes of this analysis (Supplementary Figure 7). In this case, 13 and 20 terms were retrieved for *lncRNA45*WT and *lncRNA45*MUT, respectively, with 12 terms shared between both protein lists. Consistently, RNA processing (GO:0006396) was exclusively enriched in *lncRNA45*WT.

### Complementation of Δ*lncRNA45* with *lncRNA45*WT restores promastigote growth

Since it was observed that *lncRNA45*-protein interactors are altered depending on the secondary structure formed by Locus 709, we investigated whether the biological function of *lncRNA45* is also related to this conserved secondary structure. To do this, similar to the pulldown assays, the wild-type (*lncRNA45*WT) and mutant (*lncRNA45*MUT, carrying the C50G substitution at Locus 709) sequences of *lncRNA45*, were cloned into the pSSU-Sat plasmid, the same plasmid used for overexpressing this lncRNA in OE_*lncRNA45* transfectants. After cloning, the plasmids were sequenced to ensure C50G was the only SNP present in the sequence and transfected into the *L. braziliensis* M2903 Δ*lncRNA45* cell line. The plasmid carrying the GFP gene instead of the *lncRNA45* sequence (pSSU-Mock) was also transfected into Δ*lncRNA45* as a control. Thus, three clonal lines were generated over the *L. braziliensis* M2903 Δ*lncRNA45* background (Figure 7A).

**Figure 7-.**
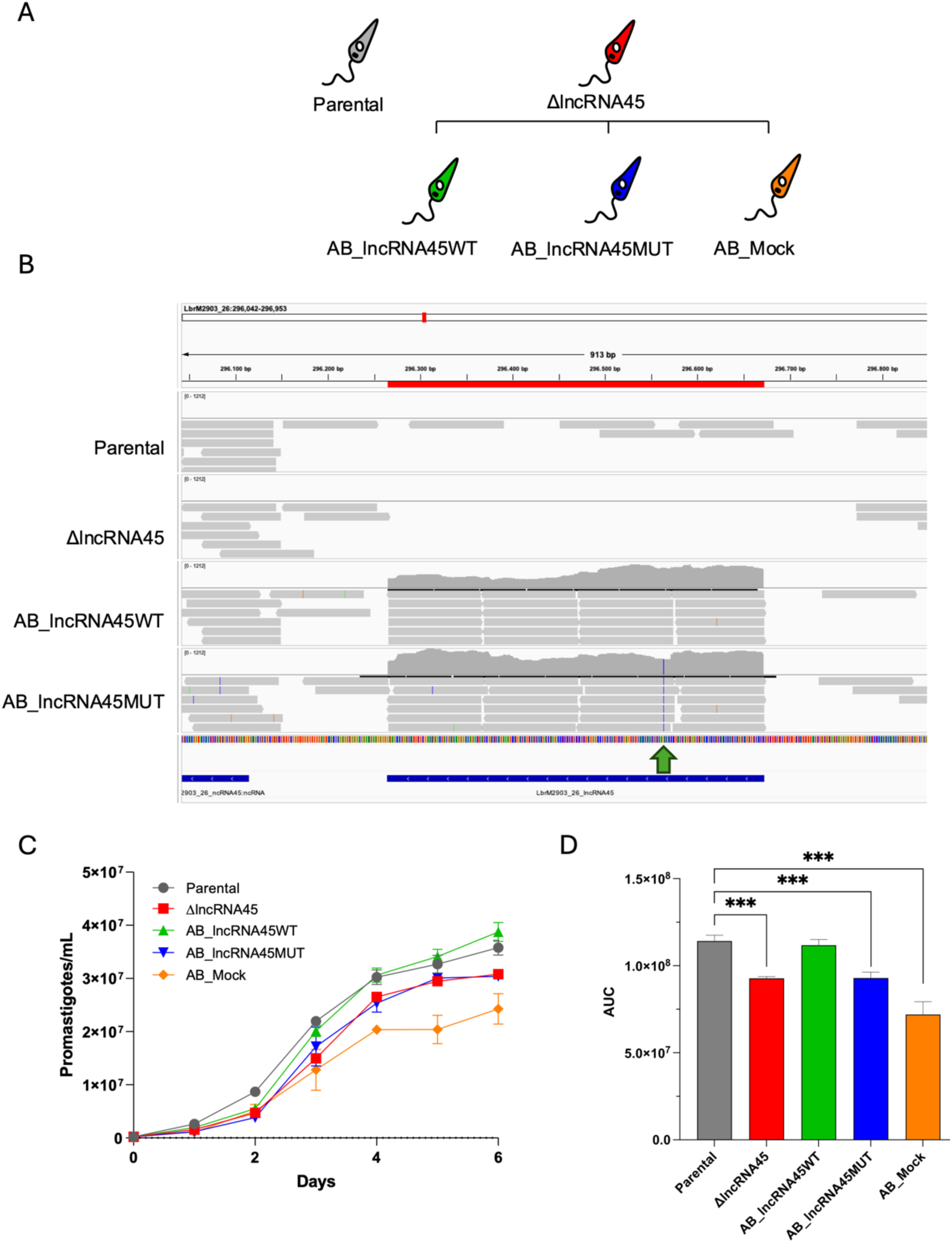
*Complementation of ΔlncRNA45 with lncRNAWT and lncRNA45MUT Sequences*. (A) The add-back cell lines of ΔlncRNA45 (red) were generated by transfecting the pSSU-SAT plasmid containing the lncRNA45WT (green) sequence, the sequence carrying the C50G mutation (lncRNA45MUT - blue), and the GFP-coding gene (AB-Mock - orange). (B) IGV representation of transcriptomic analysis of Parental, ΔlncRNA45, AB_lncRNA45WT, and AB_lncRNA45MUT cell lines. The green arrow indicates the C50G nucleotide substitution present only in AB_lncRNA45. No reads corresponding to lncRNA45 were found in the ΔlncRNA45 transcriptome, confirming the knockout. (C) Growth curves were obtained by counting culture density every 24 hours over 7 days for each transfectant and for the parental cell line (gray). (D) Significance was determined by area under the curve analysis, followed by one-way ANOVA and Tukey’s multiple comparison test.

The absence of endogenous *lncRNA45* was confirmed by PCR in all the transfectants (Supplementary Figure 6G - PCR A). The presence of the plasmid containing *lncRNA45*WT or *lncRNA45*MUT was confirmed in the add-back cell lines AB_*lncRNA45*WT and AB_*lncRNA45*MUT, respectively (Supplementary Figure 6G - PCR B). The presence of the plasmid pSSU-GFP was confirmed in the AB_Mock cell line (Supplementary Figure 6G - PCR G). Amplification of a 0.32 kb fragment of *hsp70* CDS was used as control for DNA presence and integrity (Supplementary Figure 6G - PCR D). RNA-seq data analysis also confirmed the absence of reads mapping to *lncRNA45* in the Δ*lncRNA45* cell line, proving that this lncRNA was indeed deleted (Figure 7B). The RNA Seq data further confirmed expression of *lncRNA45* was re-established in the add-back and that C50G is the only nucleotide substitution in the *lncRNA45*MUT sequence (Figure 7B – green arrow).

A 7-day growth curve was generated, as described previously. As observed before, reduced growth was seen in Δ*lncRNA45* compared to the parental cell line (Figure 5A and Figure 7C-D). The add-back of the *lncRNA45*WT sequence restored the growth of Δ*lncRNA45* to the levels of the parental cell line (Figures 7C-D). Supporting the significance of Locus 709 and requirement of *lncRNA45*, neither the *lncRNA45*MUT nor mock add-back control (AB_Mock) can restore Δ*lncRNA45* growth to the levels of the parental line levels (Figures 7C-D).

### *lncRNA45* does not regulate other mRNA levels

The finding that *lncRNA45* binds to a set of proteins involved in RNA processing machinery in its native sequence, but not in the presence of the C50G substitution (Figure 6D), supports the hypothesis that *lncRNA45* may regulate the abundance of other transcripts. To investigate this, RNA sequencing was conducted for the parental cell line, Δ*lncRNA45*, AB_*lncRNA45*WT, and AB_*lncRNA45*MUT cell lines. Only two genes were found to be differentially expressed in the knockout (Δ*lncRNA45*) compared to the parental cell line (DESeq2 p < 0.05, FC > 1.5) (Figure 8). No significant differences in gene expression were observed between the knockout and add-back cell lines, suggesting that *lncRNA45* does not regulate global mRNA levels in *L. braziliensis*.

**Figure 8-.**
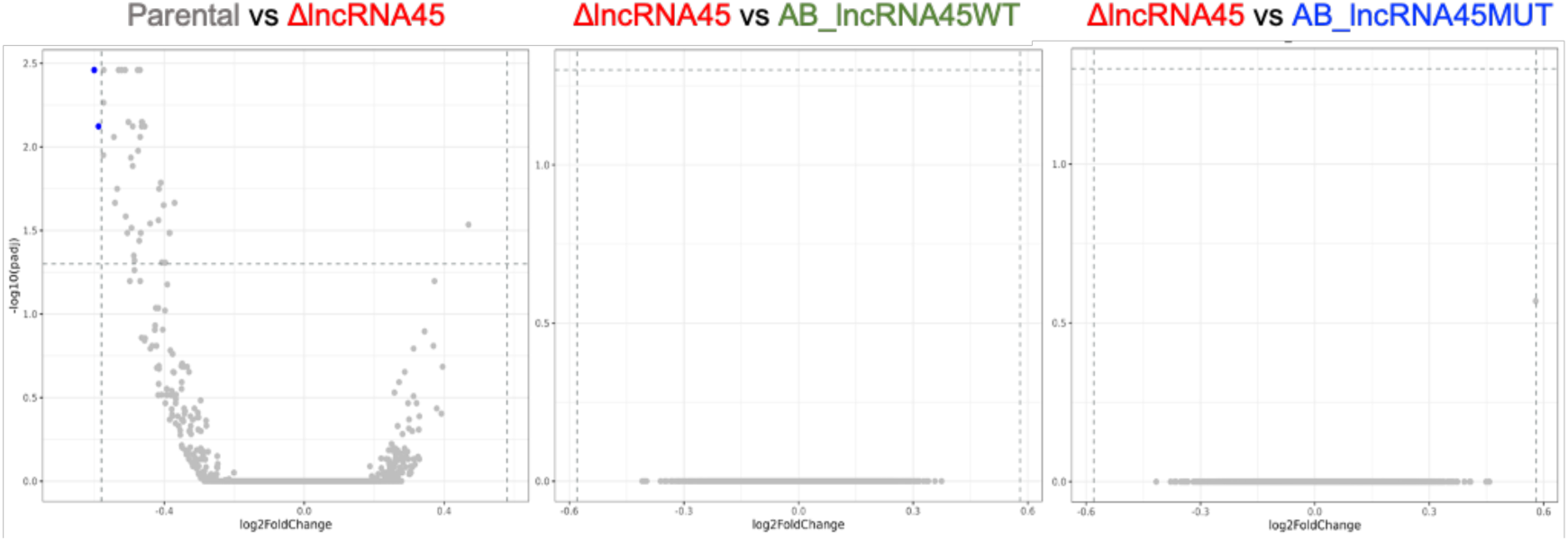
Transcriptomic analysis of lncRNA45 knockout and add-back cell lines. Volcano plots of differential expression (DESeq2, p < 0.05, FC > 1.5x) analysis between (A) Parental vs. knockout (ΔlncRNA45), (B) knockout vs. add-back complemented with the WT sequence of lncRNA45 (AB_lncRNA45WT), and (C) knockout vs. add-back complemented with the lncRNA45 sequence containing a C50G substitution at Locus 709 (AB_lncRNA45C50G). Blue dots represent genes with significant differential expression, while gray dots indicate non-significant differences.

### *lncRNA45*-protein interaction is dependent on Locus 709 secondary structure

RNA/Protein immunoprecipitation (RIP:IP) was conducted to confirm *in vivo* relevance of *lncRNA45*-protein interactions identified in the S1m *in vitro* pulldown assay. The five proteins that interact only with *lncRNA45*WT and not with *lncRNA45*MUT were tagged with HA at the N-terminus (Supplementary Figure 8) and immunoprecipitated using anti-HA magnetic beads. The *L. braziliensis* M2903 parental cell line (untagged) was used as the negative control.

RNA that immunoprecipitated with the tagged candidate interacting proteins was extracted, rRNA-cleared and sequenced to confirm interaction with *lncRNA45* and identify other potential transcripts in the complex. Surprisingly, *lncRNA45* was enriched only in the eluate of the hypothetical protein (Lb4 - LBRM2903_280022100). A higher number of counts of *lncRNA45* transcript was also observed for XRN 5’-3’ exonuclease N-terminus (Lb2 - LBRM2903_160009500) in comparison to the negative control, though this difference was not statistically significant (Figure 9).

**Figure 9-.**
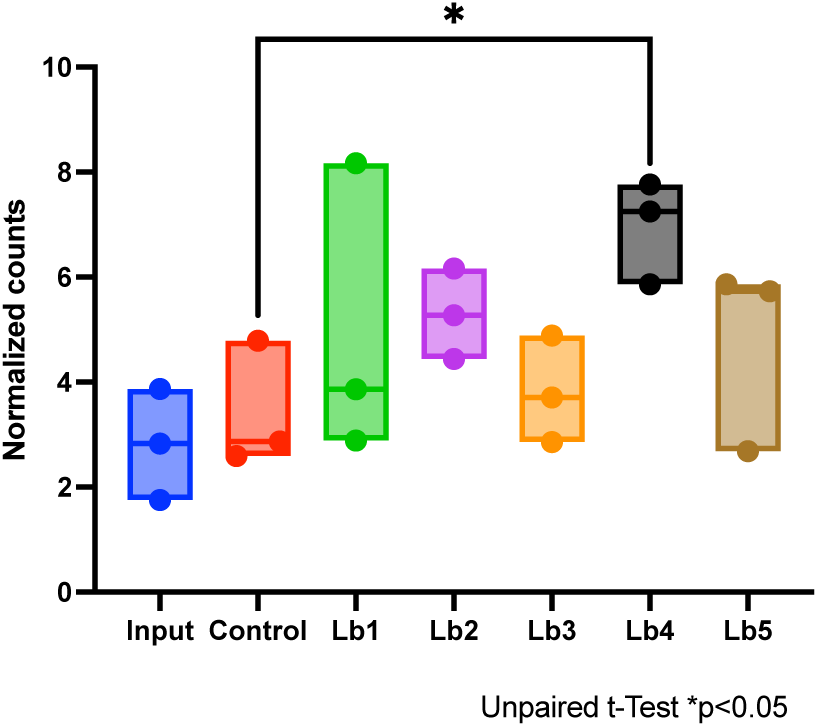
lncRNA45-protein interaction was partially confirmed by RIP. RNA immunoprecipitated with the tagged proteins (Lb1 – Lb5), with the negative control (untagged parental cell *L. braziliensis* M2903) and the total RNA of the parental cell line (Input) were extracted, treated with DNAse. Before sequencing, samples were depleted for rRNA. Raw counts of three biological replicates were normalized based on library size, plotted and tested for significance. lncRNA45 was significantly enriched only in Lb4 eluate (Unpaired t-Test *p<0.05).

In the same assay, protein-protein interactions were investigated by mass spectrometry to evaluate potential networks involving *lncRNA45*. Interestingly, four out of the five analyzed proteins were found to interact with at least one of the 18 proteins identified as *lncRNA45*WT-binding in the S1m pulldown assay (Figure 6A and 6C), confirming some of the interactions predicted by STRING (*Search Tool for the Retrieval of Interacting Genes/Proteins*) (57).

To determine if these protein-protein interactions are RNA-dependent, a fraction of the eluate material was treated with RNAse and compared with the untreated eluate. Our findings show that the interaction of Lb1 (40S ribosomal protein) with Lb3 (Lupus La protein homolog) is lost after RNase treatment (Figure 10B and 10D). The same is true for the interaction of Lb2 with Lb1, Lb3, Lb8, Lb11, and Lb12 (Figure 10B and 10D). Three other protein-protein interactions (Lb5-Lb4, Lb5-Lb6, Lb2-Lb14) that were lost in RNase-treated compared to untreated samples were not significantly different from the negative control and may thus be artifacts (Figure 10B and 10D). Notably, none of these proteins were detected in the negative controls, confirming their specific interaction with identified targets. This supports the significance of *lncRNA45* in regulatory protein complex interactions and functional cohesion.

**Figure 10-.**
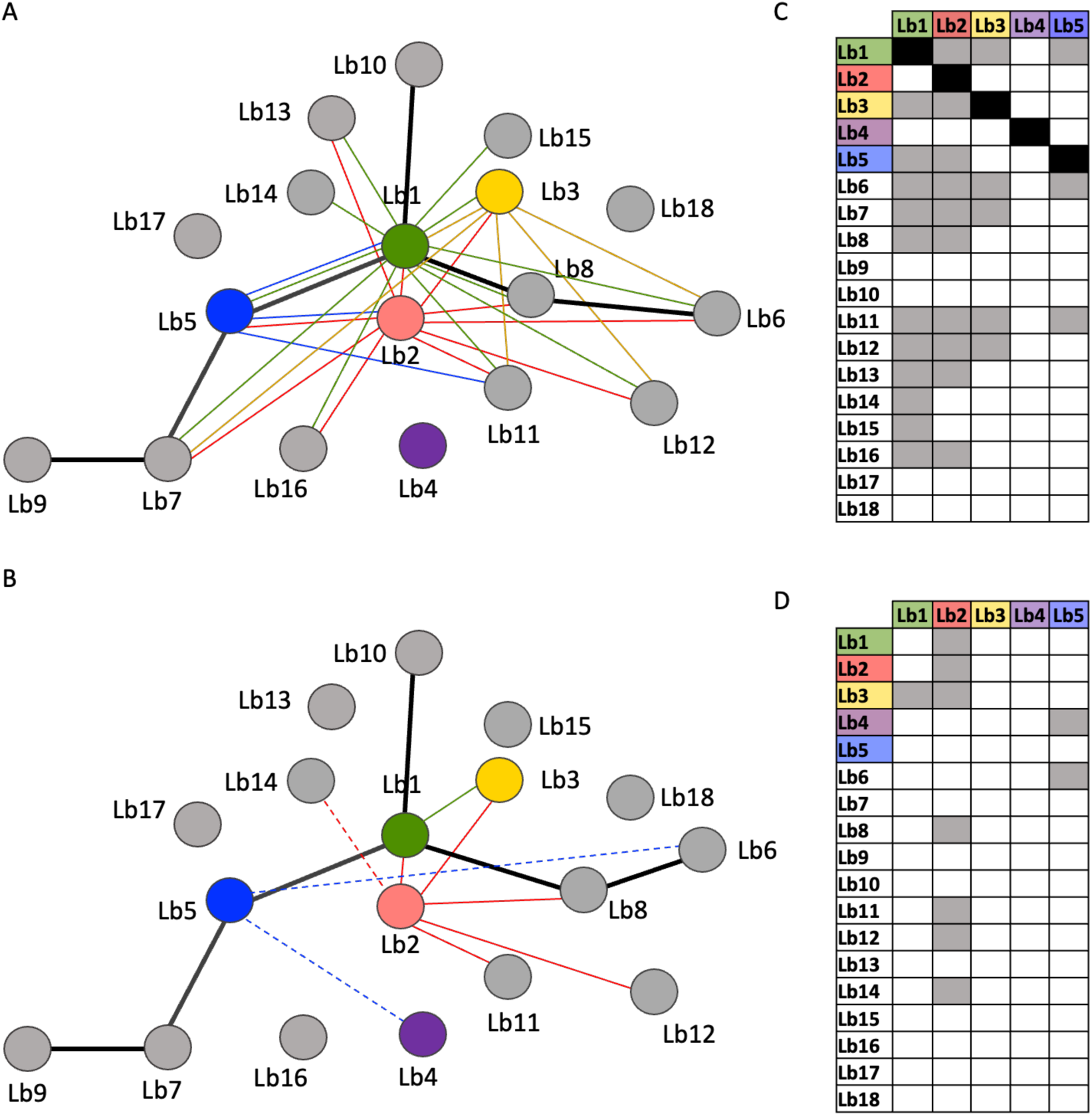
Network analysis of in silico predicted and RIP-identified interactions between proteins identified as lncRNA45 binders. (A-B) Each circle represents one of the 18 proteins identified in the S1m assay as lncRNA45 interactors, labelled Lb1-Lb18 (gene identification details are provided in Table 1). Coloured circles represent the five proteins characterized in RIP assays. Black bold lines connecting the circles indicate connections identified in silico using String. Coloured lines represent the interactions of the protein shown in the corresponding colour, empirically determined in RIP MS analysis. (A) Comparison between proteins identified in the eluate of RIP samples (HA-tagged) and the negative control (proteins found in the eluate of the parental cell line). (B) Comparison of proteins identified in the eluate of RIP samples treated or untreated with RNase. Dotted lines indicate interactions identified only in the comparison between treated and untreated samples, that is, protein-protein interactions are indirect in these cases, depending on RNA, but are absent in the comparison of target versus negative control. (C-D) Diagram of protein-protein interactions identified in RIP experiments for untreated (C) and RNAse treated samples (D). Boxes filled in Gray show identified interactions. Boxes filled in black represent the identification of the target protein in the sample.

## DISCUSSION

In this study, we used innovative *in silico* methodologies to classify putative ncRNAs of *Leishmania* based on the evolutionary conservation of their structure. Using reverse genetic tools, we demonstrate a novel ncRNA with a conserved secondary structure, *lncRNA45*, is a regulatory ncRNA impacting cellular fitness whose structure is critical to function.

Our applied computational approach combined two strategies for genome-wide screens of structured RNAs. It combined simultaneous use of sequence and structural alignment strategies and included the evolutionary conservation of secondary structure. To discover RNA families sharing the same structure, we used methods for clustering conserved RNA structures. The analysis of the structure-based RNA clusters conferred a rational basis to select ncRNAs for further investigation by reverse genetics, and we analyzed the functional impact of a SNP that modifies *lncRNA45* structure.

It is well established that gene expression modulation throughout the parasite lifecycle occurs and differences in RNA abundance between clinical isolates or under distinct environmental conditions have been demonstrated for *Leishmania* and other trypanosomatids (4,5,58,59). How these parasites modulate gene expression in a context of polycistronic transcription and lack of canonical promoters, or the factors and mechanisms used to regulate gene expression are still not completely understood (6). Only recently have some studies suggested the existence of ncRNAs in trypanosomatids. Interestingly, computational and functional assays have demonstrated that among trypanosomatids only *Trypanosoma brucei* and *Leishmania Viannia* species have functional RNAi machinery (60). But, so far functional exploration and discovery of putative regulatory ncRNAs is still incipient in *T. brucei* and *Leishmania* spp. (16,61). Nevertheless, the more recent discovery of ncRNAs in trypanosomatids (12,16–18) introduced a new factor in gene expression control, complementing the better characterized RNA-binding proteins (5,62,63). Until recently, it was unclear whether ncRNAs had biological and regulatory functions in trypanosomatids. However, in 2022, (12) showed that overexpression of *Grumpy* lncRNA leads to an increase in *T. brucei* differentiation to stumpy forms, increasing survival rates in mice infected with *T. brucei*.

It is well known that lncRNAs are poorly conserved in terms of sequence in different organisms (64). Nevertheless, it has been shown that ncRNA transcripts have specific structured regions, some even syntenically conserved between organisms, that are associated not only with stability but also with biological function (65). In an effort to search for structured regions conserved in the genome of trypanosomatids, we conducted a computational analysis to uncover secondary structure evolutionary conservation and selected one target, *lncRNA45* for functional characterization. Here, we present for the first time an *in silico* and functional characterization of a structured lncRNA in *L. braziliensis*.

*LncRNA45* was identified in a comprehensive transcriptomic study aimed at deciphering gene expression regulation mechanisms in *L. braziliensis* (17). This lncRNA was selected among others with a conserved secondary structure, because the RNAseq analysis previously conducted indicated that it was differentially expressed throughout the lifecycle and is preferentially expressed in the intracellular, proliferative form that infects mammalian macrophages (17). *LncRNA45* contains a structured motif (Locus 709), conserved in 6 *Leishmania* species (*L. braziliensis*, *L. panamensis*, *L. major* Friedlin, *L. donovani*, *L. infantum* and *L. amazonensis)* suggestive that this lncRNA structure may have a conserved essential function throughout *Leishmania*. In contrast, the full *lncRNA45* sequence is conserved only in four *Leishmania* species, all belonging to the *Viannia* subgenus. The fact that Locus 709 sequence is also conserved only in these four species,

The presence of *lncRNA45* in *L. braziliensis* was confirmed by several approaches (RNAseq, RT-qPCR, RNA FISH, circularization assay and northern blotting) showing that this transcript is consistently expressed in this parasite, and it is not an artifact of the RNASeq analysis previously conducted (17). The characterization of *lncRNA45* shows it does not undergo the same processing routes as mRNAs in *L. braziliensis*, a common feature of lncRNAs observed across other organisms (66). This appears to be a characteristic feature of ncRNAs in *Leishmania* as well (13). *LncRNA45* has a short (17nt) poly (A) tail but lacks any traits of the spliced-leader sequence in the 5’ end. The treatment with a decapping enzyme (TAP) suggests that there is a protection of the 5’end and that *lncRNA45* is not a monophosphorylated transcript. Further studies must be conducted to understand if alternative capping occurs and what is the process of biogenesis of the mature *lncRNA45* transcript and how its stability is achieved to permit its protection from degradation and functionality (67).

Loss-of-function analysis showed that *lncRNA45* has a direct role in parasite growth in culture, correlating with the increased expression of this lncRNA in this life stage. Interestingly, the reduced cell density observed in the knockout compared to the parental cell line was not caused by an increase in doubling time and could be the result of an increased death rate or an altered cell signaling pathway that could be impacting quorum sensing capacity. Since no differences in capacity to recover from nutritional stress was observed, the hypothesis that these mutants could be more susceptible to nutritional stress and thus would stop multiplying at some point was ruled out.

To test if this biological role is structure-dependent, we predicted *in silico* a single nucleotide substitution in Locus 709 (C50G) that theoretically leads to an impairment of its secondary structure. The fact that the *lncRNA45* sequence carrying the C50G modification (*lncRNA45*MUT) addback was unable to recover the growth phenotype of the knockout cell line as the line complemented with the wild-type sequence (*lncRNA45*WT) confirmed the essentiality of Locus 709 for *lncRNA45* activity. Similarly, mutations in structured lncRNAs, such as *lnc-31* and *Braveheart* lncRNA, impair their ability to sustain molecular interactions, such as scaffolding of proteins or RNA, that are critical for their function (20,68).

It is well known that the functional mechanism in which a lncRNA is involved is dependent on its subcellular location (14). Cytoplasmic lncRNAs are more commonly involved in regulating mRNA stability, RNA processing, translation rates or protein function, whereas nuclear lncRNAs were shown to act by regulating transcription itself (69). RNA FISH revealed that *lncRNA45* is a cytoplasmic lncRNA and does not present a preferential location, being spotted in different regions of *L. braziliensis*’ cytoplasm. Additionally, RNAseq showed that *lncRNA45* is not involved in the regulation of specific mRNAs in this parasite. These findings led us to hypothesize that *lncRNA45* could be acting on RNA processing, impairing or increasing, for example, translation rates; else directly regulating protein function and stability. Similar results were found for *lnc-31* regulation of *Rock1*, in which although the mRNA levels of *Rock1* remained unchanged in the cell line knockdown for *lnc-31*, protein expression was notably downregulated suggesting a potential positive role of lnc-31 in promoting the translation of Rock1 mRNA (68).

The *in vitro* pulldown assay using as bait both, *lncRNA45*WT and *lncRNA45*MUT sequences confirmed the importance of the secondary structure formed by Locus 709 for *lncRNA45* activity, showing that the presence of the C50G substitution impairs the binding of five proteins to this lncRNA. The group of proteins pulled-down with *lncRNA45* suggests this transcript may be involved in regulating RNA processing or cell-cell signaling pathways, mainly quorum sensing which corroborates the phenotype of reduced parasite density in culture observed in loss-of-function assays.

Although lncRNA-protein interaction was not statistically confirmed for all the five proteins submitted to RIP assays, our RNAseq results show a higher number of *lncRNA45* counts in the protein eluates compared to the negative control for XRN 5’-3’ exonuclease (Lb2) and two biological replicates of Nop53 (Lb5). For the hypothetical protein (Lb4), *lncRNA45* was significantly enriched in the eluate. It is worth noting that *lncRNA45* is not an abundant transcript in *L. braziliensis* M2903, as observed in the RNAseq data of the parental cell line, and it is possible that this low abundance could impair the statistical analysis, since for other more abundant lncRNAs investigated in the lab, this effect was not observed. Interestingly, interaction between Lb4 and Lb5 proteins was shown to be RNA-dependent and interaction of Lb2 with other 6 proteins (Lb1, Lb3, Lb8, Lb11, Lb12 and Lb14) was also shown to be RNA-dependent, suggesting that *lncRNA45* could be responsible for maintaining the complex together.

An in-depth literature search of the five proteins interacting with *lncRNA45* in a structure-dependent manner showed that at least one of them, Lupus La homolog was identified as an interactor of other two long non-coding RNA: XIST in mouse and Grumpy in *T. brucei*. In addition, this same protein (Lupus La homolog) and XRN 5’-3’ exonuclease were shown to directly impact *T. brucei* growth in loss-of-functions studies (70), as observed for *L. braziliensis* knockout for *lncRNA45*. It is interesting to note also that three of these proteins (40S ribosomal protein S9, Nop53 and Lupus La) were previously associated with ribosomal biogenesis and translation initiation (42–44). RIP-seq MS data not only confirmed but expanded the interaction network observed for the 18 proteins identified as *lncRNA45* interactors. Additionally, it is worth to highlight Lupus La homolog interaction with both 40S ribosomal protein and Nop53 was lost in RNAse-treated RIP samples, suggesting *lncRNA45* and/or other RNAs are necessary for these protein-protein interaction.

To our knowledge, this is the first demonstration of secondary structure-dependent lncRNA activity in trypanosomatids, as well as of a regulatory ncRNA in *Leishmania.* This study encourages the search for other regulatory conserved structured lncRNAs that could be targeted for treatment or development of vaccines.

## CONCLUSIONS

In conclusion, our study highlights the significance of long non-coding RNAs (lncRNAs) in the regulation of gene expression within trypanosomatids. By characterizing *lncRNA45* in *Leishmania braziliensis*, we demonstrated that the conserved secondary structure in this lncRNA is essential for its interaction with proteins and its biological function. Our loss-of-function analysis reveals *lncRNA45* plays a crucial role in promastigote growth, emphasizing the importance of its secondary structure for its function.

The interaction network identified through pulldown assays and RIP-seq underscores *lncRNA45*’s involvement in RNA processing and cell signaling pathways, notably quorum sensing. Furthermore, the identification of specific protein interactors, including Lupus La homolog, 40S ribosomal protein S9, and Nop53, suggests that *lncRNA45* participates in ribosomal biogenesis and translation initiation.

This study provides the first evidence of regulatory ncRNAs in *Leishmania,* underpinning the secondary structure-dependent lncRNA activity in trypanosomatids. This paves the way for further research on conserved structured lncRNAs as potential therapeutic targets or vaccine candidates. Overall, our findings contribute to a better understanding of lncRNA conservation and potential functions in trypanosomatids; illustrating regulatory complexity and impact upon parasite biology.

## SUPPLEMENTARY DATA

Supplementary Data are available

## AUTHOR CONTRIBUTIONS

C.R.E. contributed to the conceptualization, data acquisition, analysis, and interpretation, drafted the manuscript, and revised it. C.A. and R.D.M.M. were responsible for conceptualization of computational analysis, data acquisition, and analysis, and interpretation of secondary structure predictions, drafted sections of the manuscript, and revised it. J.C.Q.J. acquired data for growth curves and in vitro infections and contributed to manuscript revision. N.M.M. assisted with RNA immunoprecipitation acquisition and analysis. F.P. conducted RNA-seq analysis and revised the manuscript. L.A.O. performed RNA-seq analysis, interactive network analysis, and revised the manuscript. L.A. contributed to data acquisition for infection experiments. T.P.A.D. conducted RT-qPCR and RNA extraction experiments. A.D. carried out mass spectrometry acquisition and analysis of RNA immunoprecipitation and revised the manuscript. J.G. contributed to the conceptualization, data analysis, interpretation, and manuscript revision. P.B.W. participated in the conceptualization, data interpretation, secured funding, and reviewed the manuscript. A.K.C. led the conceptualization, supervised the study, secured funding, and wrote and revised the manuscript. All authors reviewed, edited, and approved the final version of the manuscript.

## ACKNOWLEDGEMENTS

We thank Dr. Susanne Kramer for assistance with the Northern blotting assay. Proteomic and RNA identification and analyses were conducted in the MAP and Genomics and Bioinformatics Labs, respectively, at the Bioscience Technology Facility, University of York (York, United Kingdom). We acknowledge Viviane Ambrosio for technical support in the laboratory. We also thank Roberta R. Costas Rosales and Elizabete Rosa Milani from the Microscopy Facility at the Ribeirão Preto Medical School (Ribeirão Preto, Brazil). We appreciate Prof. Michael Plevin’s input on the *Leishmania* in-vitro pull-down experiments. Finally, we thank the staff of the Proteomics Platform at the CHU de Québec Research Centre (Quebec, Canada) and the TriTrypDB database.

