## Supplementary figures for "Functional Characterization of a Novel Long Non-Coding RNA in *Leishmania braziliensis* Identified Through Computational Screening for Conserved RNA Structures"

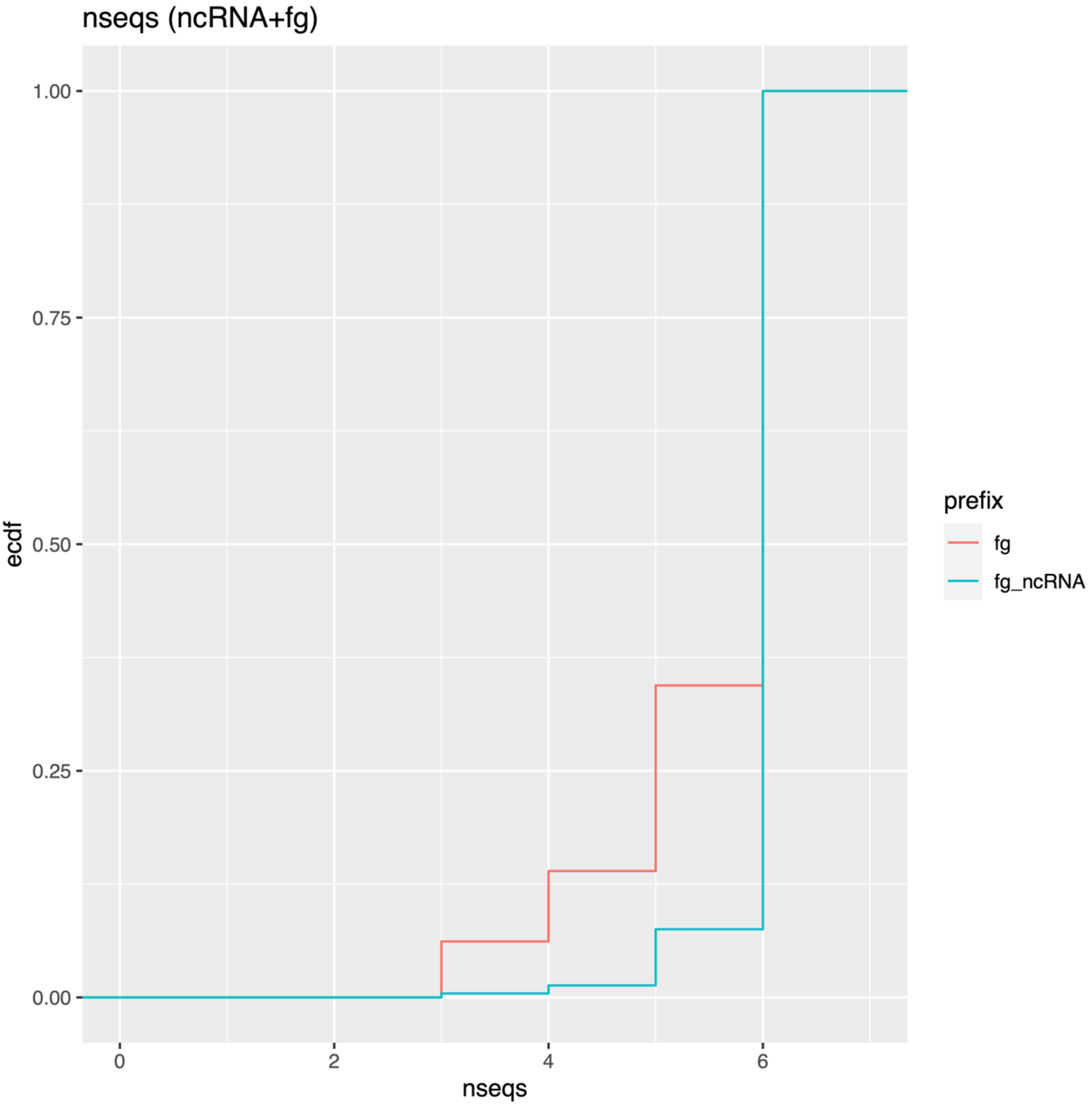


Supplementary Figure S1 - Number alignment windows with >= nseqs sequences in the 631 alignment windows predicted to be ncRNAs by RNAz (SVM score >=0) (red, fg) and in the subset of these which overlap known ncRNAs (blue, fg_ncRNA). Based on this we filter the ncRNA predictions of interest down to those with >= 6 sequences in the alignment and loose only a small fraction of the known ncRNAs.

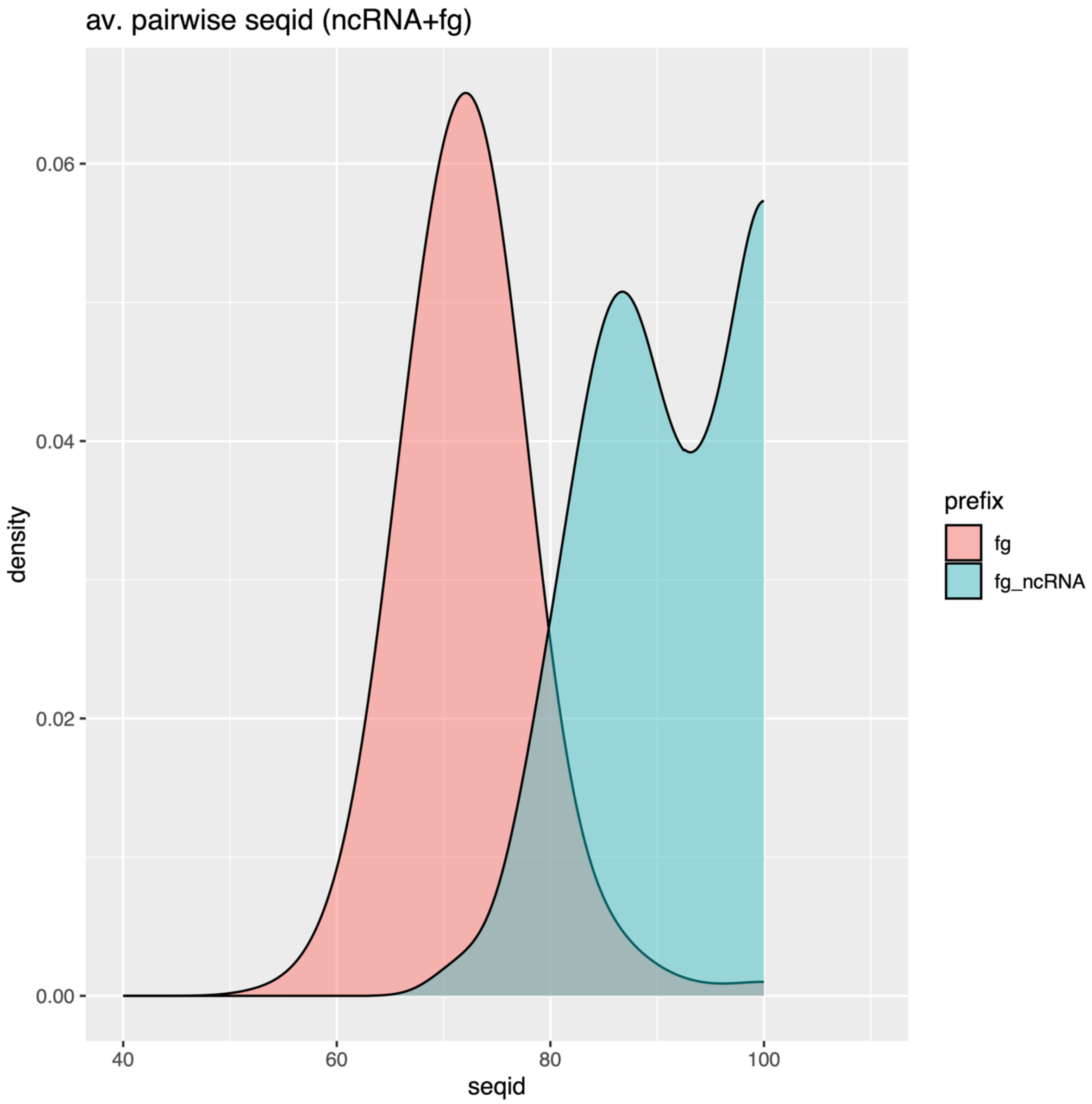


Supplementary Figure S2 - Density plot of average pairwise sequence id in the 631 alignment windows predicted to be ncRNAs by RNAz (SVM score >=0) (red, fg) and in the subset of these which overlap known ncRNAs (blue, fg_ncRNA). Based on this we filter the ncRNA predictions of interest down to those sequence id >= 70% and loose only a small fraction of the known ncRNAs.

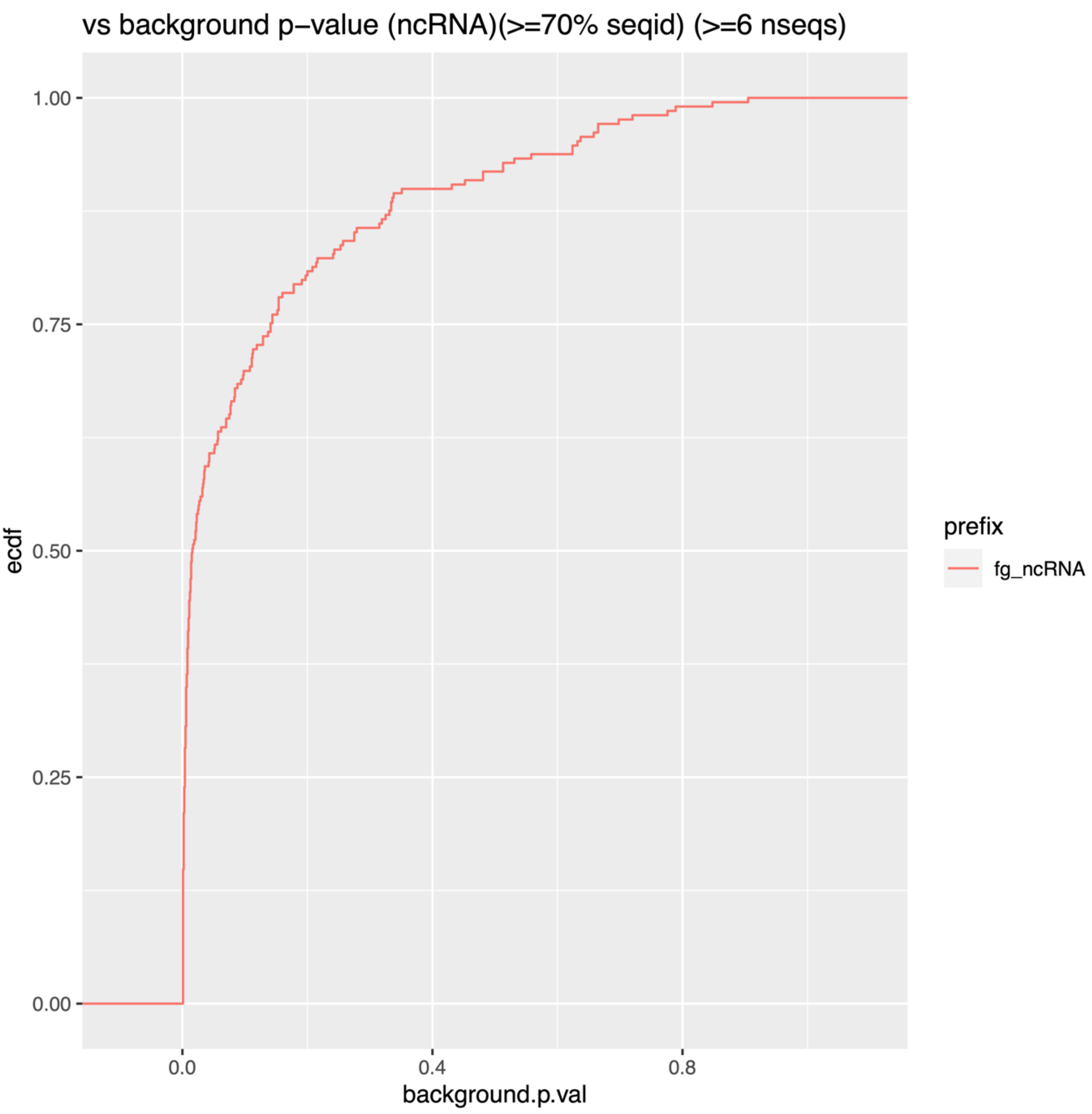


Supplementary Figure S3 - p value for the subset of the known ncRNAs which have sequence id >= 70 and at least 6 sequences in the alignment window. The background is 100 random shufflings with sissiz of the 12,969 alignment windows. The chosen cutoff based on this plot and on the details in the supplementary RNAz table is p<=0.004.

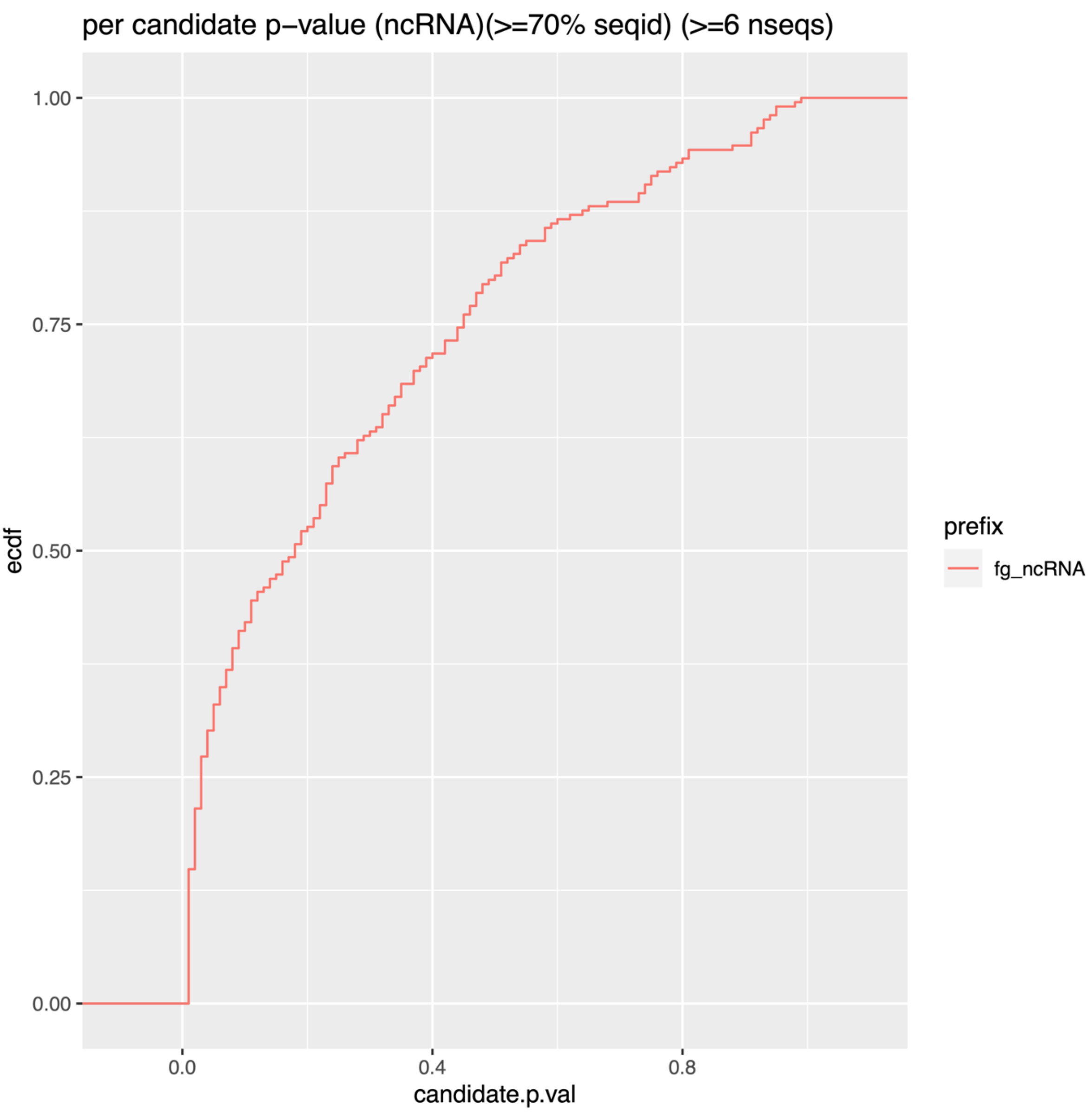


Supplementary Figure S4 - per candidate p value for the subset of the known ncRNAs which have sequence id >= 70 and at least 6 sequences in the alignment window. The background for a particular candide is 100 random shufflings with sissiz of the candidates own alignment window. The cutoff is conservatively chosen as the minimum (1%).

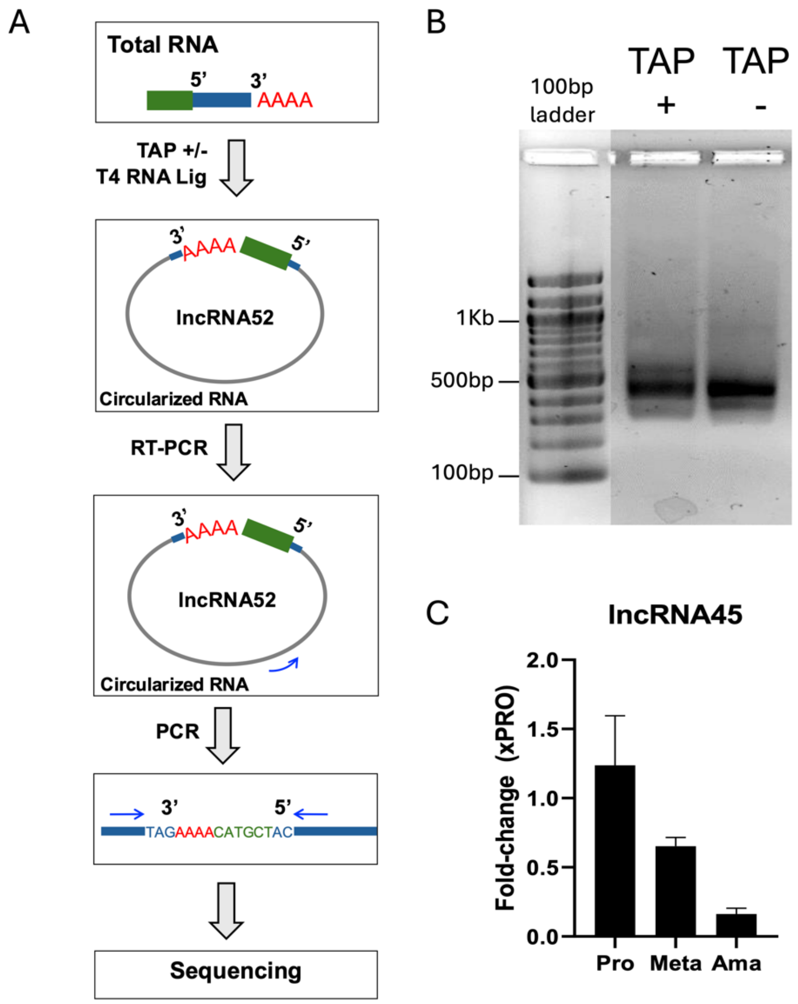


Supplementary Figure S5 – (A) A protocol of RNA circularization was used to determine the lncRNA size. In this protocol, TAP-treated (TAP+) and untreated (TAP-) total RNA is circularized and reversed-transcribed using primers specific to the target. The resulting cDNA is then PCR-amplified using primers directed to the 5' and 3' ends of the transcript and submitted to sequencing. (B) Electrophoresis of the PCR from circularized RNAs treated (TAP+) or not (TAP-) with tobacco acidic phosphatase (TAP) for cap removal. Amplification was done using the cDNA obtained from circularized RNA and primers directed towards the transcript ends. A similar band was observed in both conditions. (C) Differential expression of lncRNA45 assessed by RT-qPCR using total RNAs of the three main morphologies. The endogenous gene 7SL was used for normalization.

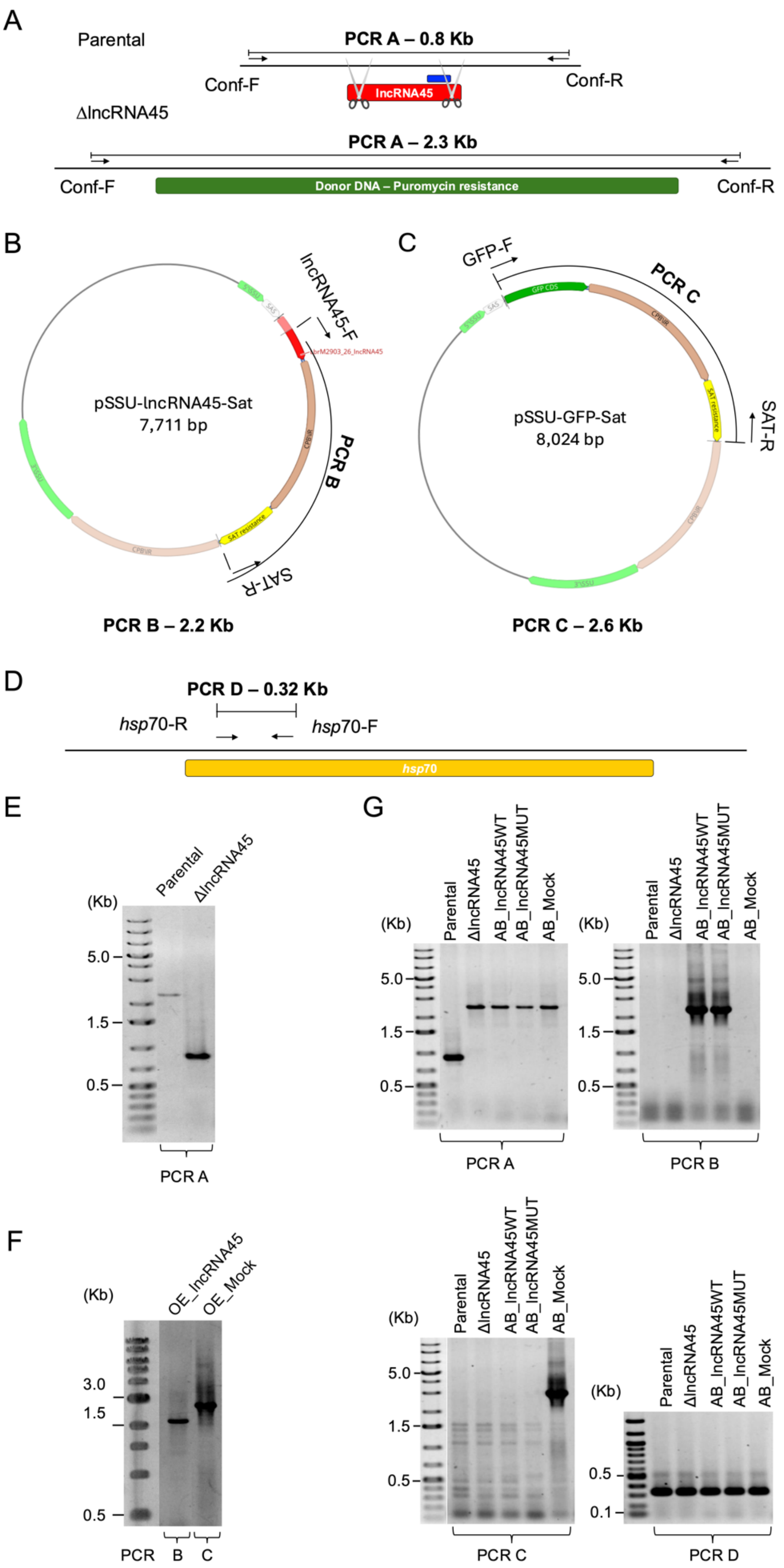


Supplementary Figure S6 – Genotypic characterization of the modified cell lines generated in this study. (A) PCR A was used to confirm the knockout of lncRNA. Using the primers Conf-F and Conf-R, annealing outside the double strand break region, a 0.8 Kb fragment was amplified in the parental line whereas in the knockout cell line, a 2.3 Kb fragment was amplified due to the insertion of the donor DNA cassette. (B)

Primers lncRNA45-F (annealing) inside lncRNA45 and SAT-R (annealing inside nourseothricin resistance gene SAT) were used to confirm the presence of the vector used for both lncRNA45 overexpression in the parental cell line, and complementation in ΔlncRNA45 cell line. (C) Primers GFP-F and SAT-R were used to confirm the presence of the mock plasmid transfected in parental (OE_Mock) and ΔlncRNA45 (AB_Mock) as a negative control. (D) Two primers annealing in *heat shock protein 70* CDS (*hsp*70-R and *hsp*70-F) were used to confirm the presence of DNA in all samples. (E) The complete knockout of lncRNA45 was confirmed by PCR A, resulting, as expected, in a 0.8 Kb and a 2.3 Kb fragments for parental and ΔlncRNA45 cell lines, respectively. (F) The presence of pSSU-lncRNA45-Sat was confirmed for the cell line overexpressing lncRNA45 (OE_lncRNA45) by PCR B. Also, the presence of pSSU-GFP-Sat was confirmed for the control cell line (OE_Mock) by PCR C. (G) Different add-back cell lines were generated to investigate the relevance of secondary structure for lncRNA45 function. PCR A shows that only in the parental cell line a 0.8 Kb fragment was detected (lncRNA45 presence), whereas for all the other transfectants, a 2.3 Kb band was amplified confirming lncRNA45 was absent. PCR B confirmed the presence of the add-back plasmid containing the native sequence of lncRNA45 in AB_lncRNA45WT cell line and the presence of the add-back plasmid containing the sequence caring the C50G substitution in AB_lncRNA45MUT cell line. PCR C was employed to confirm the presence of pSSU-GFP-Sat plasmid in the AB_Mock cell line. PCR D was employed as a positive control to confirm DNA presence and integrity.

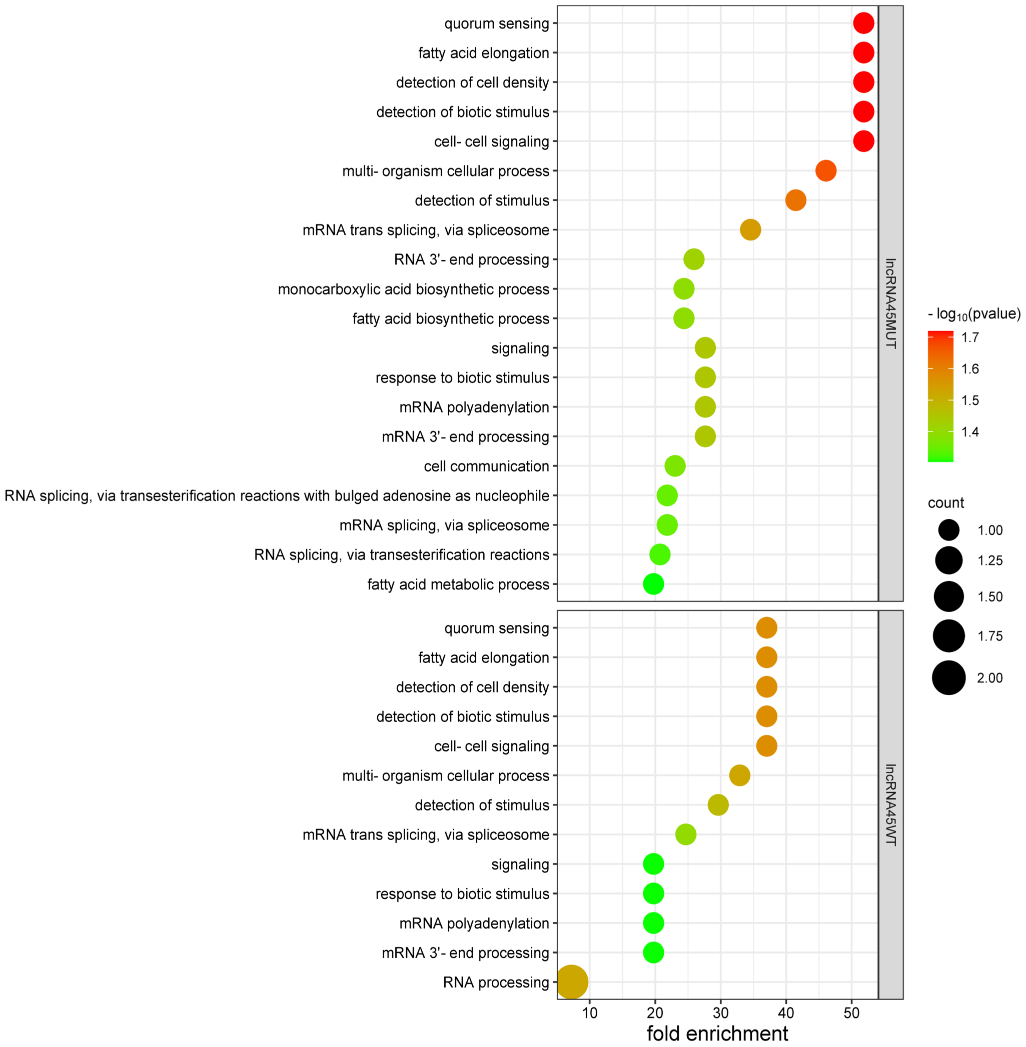


Supplementary Figure S7 - Gene ontology (GO) enrichment analysis of proteins binding to lncRNA45WT and lncRNA45MUT was performed also in *L. major* to minimize the influence of genome annotation in the analysis, since *L. major* Friedlin strain has the most complete genome annotation of *Leishmania* spp. The list of gene IDs of *L. braziliensis* M2903 and M2904 was converted into *L. major* Friedlin gene IDs and then submitted to GO enrichment analysis of biological processes in TriTrypDB. A p-value cutoff of 0.05 was used. The resulting lists of enriched terms were classified based on significance (p-value) and plotted in bubble graphs using SRPlot (http://www.bioinformatics.com.cn/srplot). The circle size (count)

represents the number of genes of a term present in the protein list. The X-axis contains the fold enrichment, which is the percentage of genes in the list relative to the percentage of genes having this term in the background. The color scale represents the significance of the enrichment determined by Fisher's exact test.

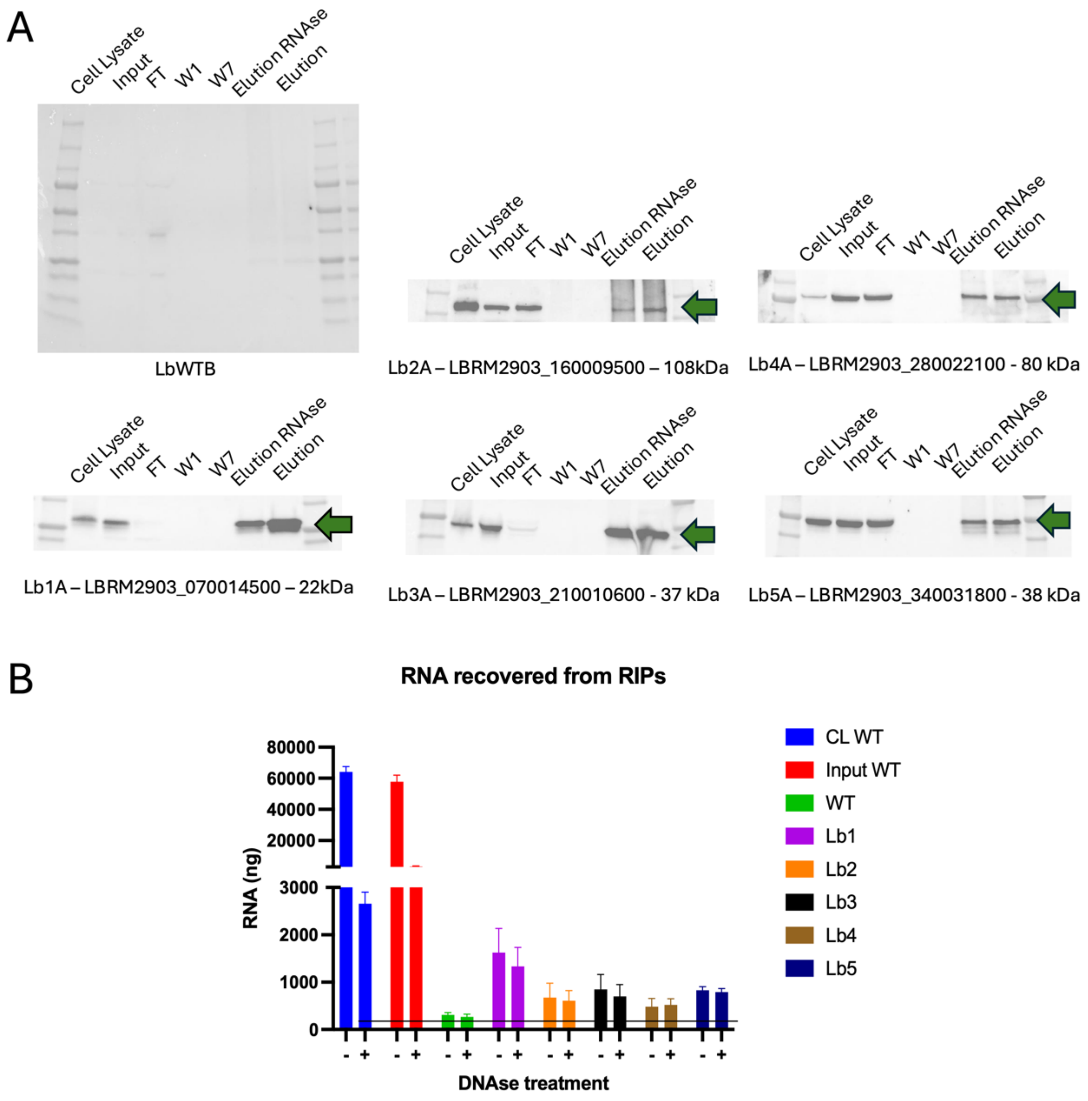


Supplementary Supplementary Figure S8 - RNA Immunoprecipitation of proteins identified as potential lncRNAs interactors in S1m pull-down. (A) Five (Lb1, Lb2, Lb3, Lb4 and Lb5) were selected for RIP assays based on the S1m pull-down results. Immunoprecipitation was done using anti-HA magnetic beads and confirmed by western blotting using anti-HA antibody. Each membrane contains the total cell lysate, the input material (cell lysate after centrifugation for debris removal), the flow through (FT), the wash 1 (W1) and wash 7 (W7) and the eluates from the samples treated (Elution RNAse) or not (Elution) with RNAse. The green arrow indicates the expected band for the tagged protein of interest. The membrane of the negative control (parental cell line LbWTB) does not present any significant signal as expected. (B) Nanodrop quantification of the RNAs recovered from the RIP assay bound to the tagged protein of interest and from the cell lysate and input of the negative control (parental cell line – WT).
